# Obtaining Spatially Resolved Tumor Purity Maps Using Deep Multiple Instance Learning In A Pan-cancer Study

**DOI:** 10.1101/2021.07.08.451443

**Authors:** Mustafa Umit Oner, Jianbin Chen, Egor Revkov, Anne James, Seow Ye Heng, Arife Neslihan Kaya, Jacob Josiah Santiago Alvarez, Angela Takano, Xin Min Cheng, Tony Kiat Hon Lim, Daniel Shao Weng Tan, Weiwei Zhai, Anders Jacobsen Skanderup, Wing-Kin Sung, Hwee Kuan Lee

## Abstract

Tumor purity is the proportion of cancer cells in the tumor tissue. An accurate tumor purity estimation is crucial for accurate pathologic evaluation and for sample selection to minimize normal cell contamination in high throughput genomic analysis. We developed a novel deep multiple instance learning model predicting tumor purity from H&E stained digital histopathology slides. Our model successfully predicted tumor purity from slides of fresh-frozen sections in eight different TCGA cohorts and formalin-fixed paraffin-embedded sections in a local Singapore cohort. The predictions were highly consistent with genomic tumor purity values, which were inferred from genomic data and accepted as the golden standard. Besides, we obtained spatially resolved tumor purity maps and showed that tumor purity varies spatially within a sample. Our analyses on tumor purity maps also suggested that pathologists might have chosen high tumor content regions inside the slides during tumor purity estimation in the TCGA cohorts, which resulted in higher values than genomic tumor purity values. In short, our model can be utilized for high throughput sample selection for genomic analysis, which will help reduce pathologists’ workload and decrease inter-observer variability. Moreover, spatial tumor purity maps can help better understand the tumor microenvironment as a key determinant in tumor formation and therapeutic response.

---

High throughput genomic analysis is indispensable for cancer research, and it has penetrated clinical practice with the promise of personalized medicine [1, 2]. One of the crucial factors affecting the quality of genomic analysis is the tumor content of the samples, which is quantified as tumor purity. A tumor consists of a complex mixture of cells, such as cancer cells, normal epithelial cells, stromal cells, and infiltrating immune cells [3], and the percentage of cancer cells within the tumor is called *tumor purity* [4].

The tumor purity affects both high throughput data acquisition and analysis. To detect genetic variations of a tumor sample by next-generation sequencing, the sample needs to have enough tumor content [5–7]. Therefore, an accurate tumor purity estimation is of great clinical importance. A sample selected based on an overestimated tumor purity, for example, may lead to a false-negative test result, which may withhold a patient from getting highly promising therapies [5]. Besides, the genomic analysis should incorporate the tumor purity to account for normal cell contamination, which can directly confound results and subsequent clinical decisions [4, 8–13]. A novel immunotherapy gene signature missed by traditional methods, for example, was discovered using a differential expression analysis incorporating tumor purity [4]. The tumor purity is also associated with clinical variables [14–16]. Low tumor purity, for instance, was associated with poor prognosis in glioma [14], colon cancer [15], and gastric cancer [16]. Moreover, tumor purity was a promising predictor for therapeutic response in colon cancer [15] and gastric cancer [16].

Tumor purity is estimated by two main approaches: percent tumor nuclei estimation and genomic tumor purity inference. A pathologist estimates tumor purity by reading H&E stained histopathology slides. Essentially, the pathologist counts the percentage of tumor nuclei over a region of interest in the slide. The tumor purity estimated in this way is referred to as *percent tumor nuclei* in this study. The percent tumor nuclei estimates are usually used for sample selection and interpretation of results in the molecular analysis. The pathologist can read any H&E stained slide and estimate percent tumor nuclei based on a cellular level analysis. Thus, this approach is widely applicable, and it has a cellular level resolution. However, counting tumor nuclei is tedious and time-consuming. More importantly, there exists inter-observer variability between pathologists’ estimates [5, 17].

Recently, tumor purity is inferred from different types of genomic data, such as somatic copy number data [18–22], somatic mutations data [23–27], gene expression data [28, 29], and DNA methylation data [30–33]. The tumor purity obtained from these methods will be referred to as *genomic tumor purity* in this study. Genomic tumor purity values are usually used in genomics analysis to mitigate confounding effects of normal cell contamination [34–36] and in correlational studies to investigate the associations between tumor purity and clinical variables [37]. Nowadays, genomic tumor purity is accepted as the golden standard. Genomic methods produce consistent values on different cancer data sets in The Cancer Genome Atlas (TCGA) [4]. However, they do not apply to the low tumor content samples. Besides, they do not provide spatial information of the locations of the cancer cells. In other words, we lose information about the spatial organization of the tumor microenvironment, which is an essential factor in therapeutic response [38]. Hence, both genomics methods and pathologists’ slide reading approach have different strengths and limitations.

This study develops a machine learning model that predicts the tumor purity from H&E stained histopathology slides such that the predictions are consistent with the genomic tumor purity values. Our model is cost-effective compared to genomics methods or pathologists’ readings since it uses readily available histopathology slides in the clinic and involves few manual steps. It also provides information about the spatial organization of the tumor microenvironment. Furthermore, previous studies showed that percent tumor nuclei estimates by different pathologists are not only inconsistent but also different from genomic tumor purity values [4, 12]. However, we still do not know the causes of the difference. This study provides some insights into the probable causes of the difference.

Two types of machine learning models can be utilized to predict tumor purity from digital histopathology slides: patch-based models and multiple instance learning (MIL) models. A patch-based model is trained on a patch cropped from a slide using the corresponding patch label determined based on pathologists’ pixel-level annotations. During inference, the predictions for all patches within the slides are obtained from the trained model, and they are aggregated to obtain sample-level tumor purity prediction. Although different studies employed this approach for tumor purity prediction [39–44], they had limited coverage since they required pathologists’ pixel-level annotations, which are rarely available, expensive, and tedious. On the other hand, the MIL paradigm does not require pixel-level annotations. It represents a sample as a bag of patches cropped from the sample’s slides and uses a sample-level label as the bag label [45–48]. Sample-level labels are weak labels providing only aggregate information rather than pixel-level information. Yet, they can easily be collected from pathology reports, electronic health records, or different data modalities.

This study designed a novel MIL model to predict tumor purity from H&E stained histopathology slides (Figure 1a). We represent each sample as a bag of patches cropped from the sample’s top and bottom slides and use the sample’s genomic tumor purity as the bag label (Methods). Our MIL model has a novel ‘distribution’ pooling filter that produces stronger bag-level representations from patches’ features than standard pooling filters like max and mean pooling (Methods). Our analysis used data from ten different TCGA cohorts and a local Singapore cohort (Table 1). The histopathology slides in each cohort were randomly segregated at the patient level into training, validation, and test sets. Then, we trained our MIL model on the training set, chose the best set of model weights based on validation set performance, and evaluated the best model on the held-out test set.

**Figure 1:**
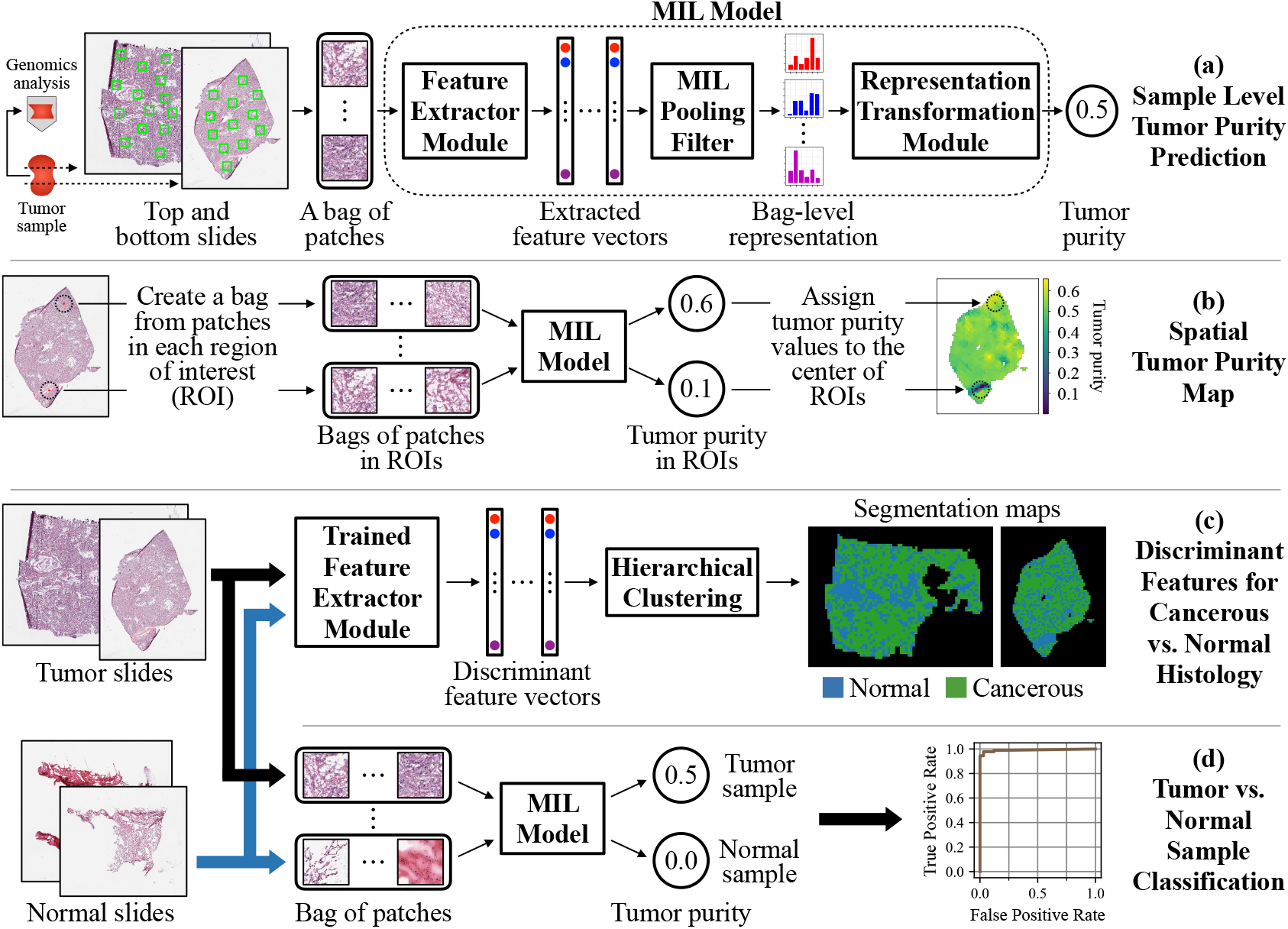
A novel MIL model predicts sample-level tumor purity from H&E stained digital histopathology slides. (**a**) Our model accepts a bag of patches cropped from the top and bottom slides of a sample as input and predicts the sample’s tumor purity at its output. The *feature extractor* module extracts a feature vector for each patch inside the bag. The *MIL pooling filter*, namely ‘distribution’ pooling, summarizes extracted features into a bag-level representation by estimating marginal feature distributions. Finally, the *bag-level representation transformation* module predicts the sample-level tumor purity. We use tumor purity values inferred from genomic sequencing data by ABSOLUTE [18] as ground-truth labels during training. (**b**) We obtain a spatial tumor purity map for a slide by inferring tumor purity over each 1*mm*^2^ region of interest within the slide in a sliding window fashion. The map shows the variation of tumor purity over the slide. (**c**) Our MIL model learned discriminant features for cancerous vs. normal histology from sample-level genomic tumor purity labels without requiring exhaustive annotations from pathologists. We used discriminant features to obtain cancerous vs. normal segmentation maps for tumor slides. Trained *feature extractor* module extracts features of patches from tumor and normal slides of a patient. Then, segmentation maps are obtained by hierarchical clustering over the extracted feature vectors. (**d**) As an essential property of tumor purity predictors, our MIL model successfully classifies samples into tumor vs. normal.

**Table 1:** The TCGA and Singapore cohorts. In each cohort, a patient has only one tumor sample and one matching normal sample if available. The number of tumor and matching normal samples in training, validation, and test sets are presented for each cohort.

| Cohorts | tumor samples |  |  | normal samples |  |  |
| --- | --- | --- | --- | --- | --- | --- |
|  | train | validation | test | train | validation | test |
| BRCA - Breast Invasive Carcinoma | 559 | 185 | 185 | 76 | 27 | 30 |
| GBM - Glioblastoma Multiforme | 285 | 95 | 94 | 0 | 0 | 0 |
| KIRC - Kidney Renal Clear Cell Carcinoma | 261 | 85 | 89 | 220 | 71 | 73 |
| LGG - Brain Lower Grade Glioma | 273 | 91 | 90 | 0 | 0 | 0 |
| LUAD - Lung Adenocarcinoma | 266 | 90 | 90 | 101 | 37 | 33 |
| LUSC - Lung Squamous Cell Carcinoma | 273 | 90 | 90 | 132 | 41 | 47 |
| OV - Ovarian Serous Cystadenocarcinoma | 310 | 103 | 103 | 53 | 13 | 18 |
| PRAD - Prostate Adenocarcinoma | 258 | 85 | 85 | 72 | 15 | 24 |
| THCA - Thyroid Carcinoma | 258 | 85 | 85 | 48 | 18 | 17 |
| UCEC - Uterine Corpus Endometrial Carcinoma | 270 | 90 | 89 | 18 | 4 | 10 |
| LUAD_SG - Lung Adenocarcinoma (Singapore) | 107 | 36 | 36 | 0 | 0 | 0 |

Our MIL models successfully predicted tumor purity from histopathology slides in different TCGA cohorts (Figure 2). The slides were of fresh-frozen sections in TCGA cohorts. However, we also showed that our MIL model could successfully predict tumor purity from H&E stained histopathology slides of formalin-fixed paraffin-embedded (ffpe) sections in the Singapore cohort using transfer learning (Figure 2i). The predictions were consistent with genomic tumor purity values. Besides, we found that the top and bottom slides of a sample were significantly different in tumor purity, which showed that tumor purity varies spatially within the sample (Figure 3c). Our findings also suggested that it was better to use both slides of the sample for tumor purity prediction whenever available. Moreover, we obtained spatially resolved tumor purity maps showing the variation of tumor purity over a slide (Figure 1b, Figure 4b, and Figure 4e).

**Figure 2:**
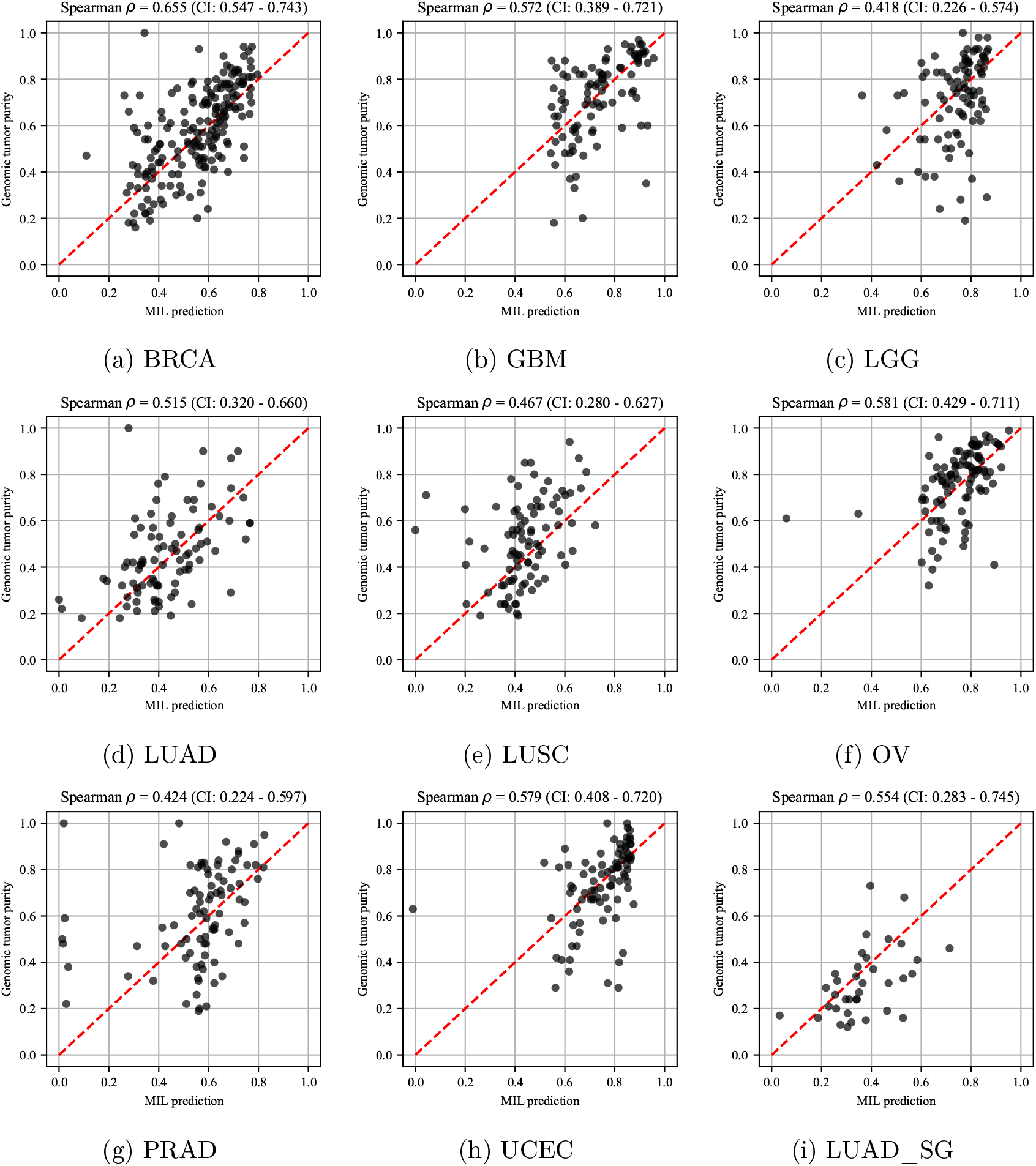
The MIL model’s tumor purity predictions correlate significantly with genomic tumor purity values. A scatter plot of genomic tumor purity obtained from ABSOLUTE [18] vs. tumor purity prediction obtained from the MIL model is given for only tumor samples in the test set of each cohort. Spearman’s correlation coefficients with 95% confidence intervals are summarized at the top of each plot. Note that the red dotted line in each plot shows the diagonal (i.e., y=x line). All data points would align on the diagonal line in case of zero prediction error.

**Figure 3:**
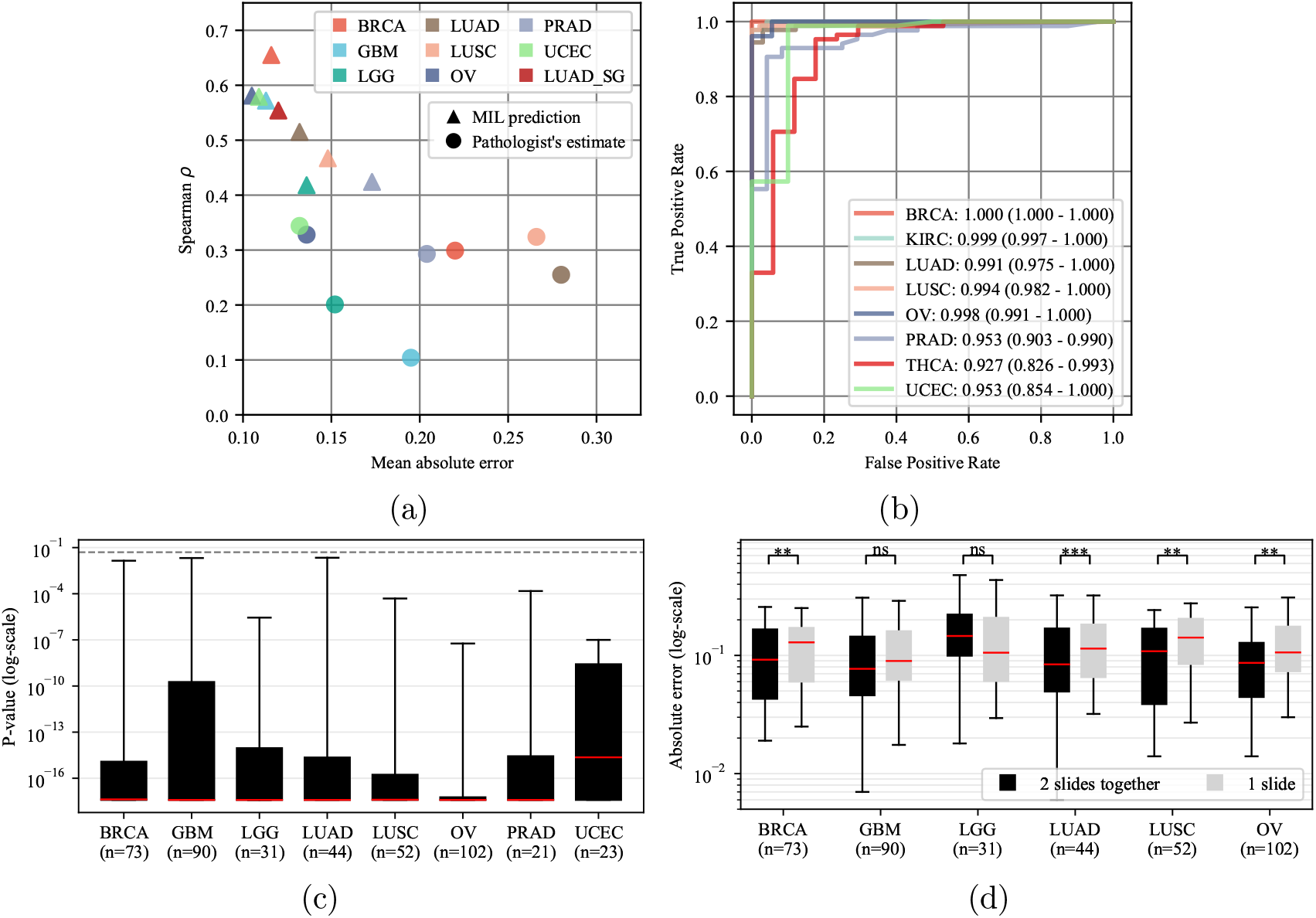
(a,b) MIL models perform better than percent tumor nuclei estimates and successfully classify samples into tumor vs. normal. (**a**) Spearman’s correlation coefficient vs. mean absolute error plot is given for MIL models’ tumor purity predictions (represented by triangles) and pathologists’ percent tumor nuclei estimates (represented by circles) in the test sets of different cohorts (showed in different colors). MIL models’ predictions achieve lower mean absolute error and higher Spearman’s correlation coefficient than percent tumor nuclei estimates. (**b**) Receiver operating characteristic curve analysis over MIL models’ predictions for tumor vs. normal sample classification. The area under curve values with 95% confidence intervals are given in the legend. MIL models successfully classified samples into tumor vs. normal in all cohorts. **(c,d) The top and bottom slides of a tumor sample are different in tumor purity.** In the test set of each cohort, for a tumor sample having top and bottom slides, we conducted two experiments. (**c**) The trained MIL model’s predictions from the top and bottom slides of a sample are statistically compared using Wilcoxon signed-rank test [52]. Each box plot summarizes the p-values obtained in a cohort. For at least 95% of the samples in each cohort, the top and bottom slides are significantly different (P<0.05) in tumor purity. The dashed line shows P=0.05. (**d**) For each sample, the absolute error between genomic tumor purity value and the MIL model’s prediction using both slides and the expected value of absolute errors between genomic tumor purity value and the MIL model’s predictions over individual slides are calculated. Box plots summarize the absolute errors in two approaches. They are statistically compared using Wilcoxon signed-rank test [52], and the results are presented on top of the plots such that *P* > 0.05 (ns: not significant), *P* ≤ 0.05 (*), *P* ≤ 0.01 (**), and *P* ≤ 0.001 (***). Whiskers show 5^*th*^ and 95^*th*^ percentiles, and red lines show median values. n: number of tumor samples with two slides.

**Figure 4:**
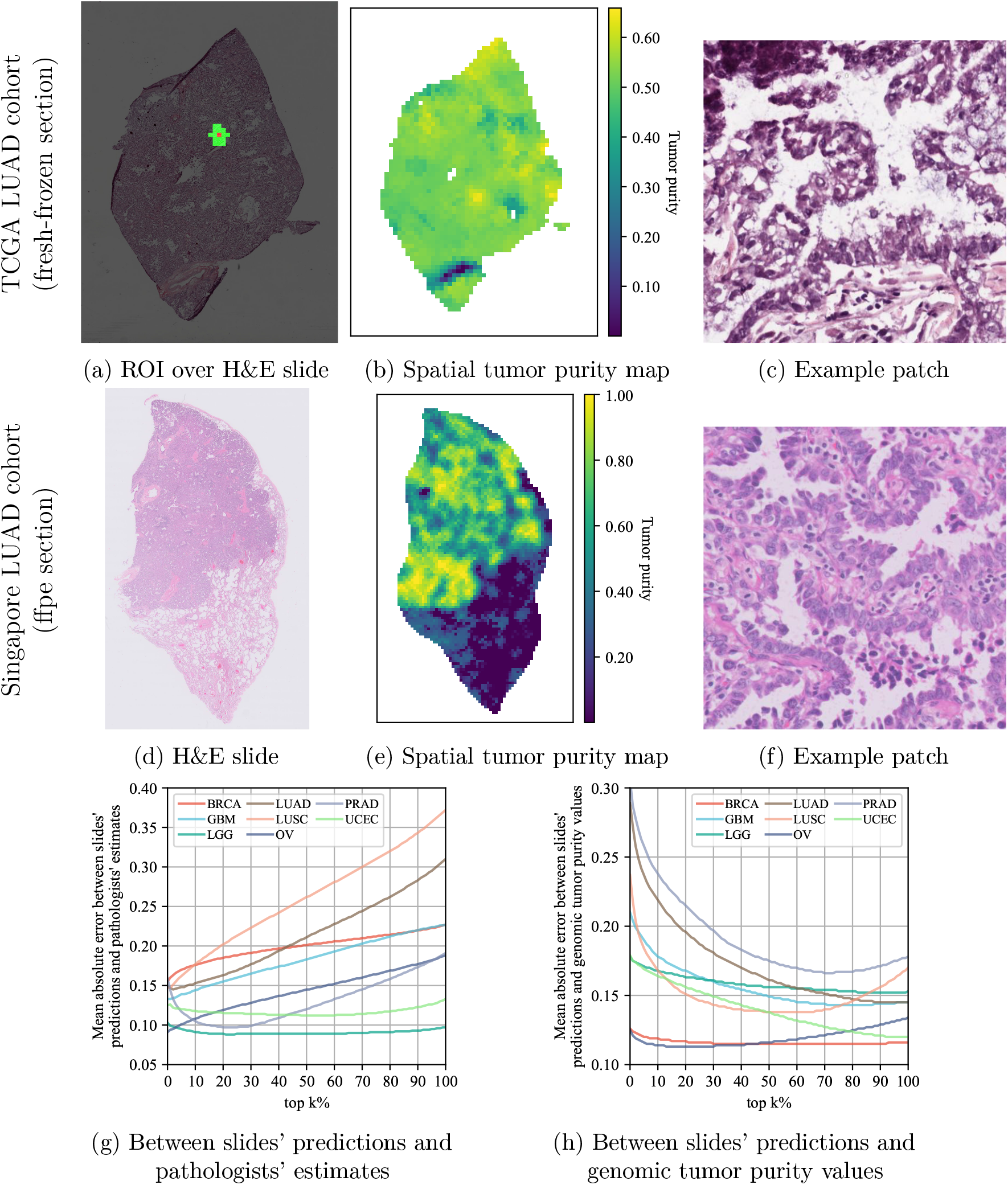
Incorrect size and selection of region-of-interest might cause overestimation in percent tumor nuclei estimates. For a slide of a fresh-frozen section in the TCGA LUAD cohort, (**a**) shows the region-of-interest (ROI) centered around a patch and consisting of 16 closest patches to that particular patch (≈ 1*mm*^2^ at the specimen level). Tumor purity corresponding to the patch is predicted over the ROI. (**b**) shows the tumor purity map for all patches within the slide. Similarly, (**d**) shows a slide of a formalin-fixed paraffin-embedded (ffpe) section in the Singapore LUAD cohort, and (**e**) shows its corresponding tumor purity map. (**c**) and (**f**) show example patches cropped from cancerous regions in the slides shown in (a) and (d), respectively. (**g, h**) To investigate the effect of the size and selection of region-of-interest on pathologists’ percent tumor nuclei estimates, we conducted error analyses over the slides’ tumor purity values by gradually extending the region-of-interest. We calculated the slide’s tumor purity as the average of top-k% of the patches with the highest scores (*k* = 0, …, 100) in the tumor purity map (*k* = 0: the patch with the highest tumor purity). In different cohorts, we plotted mean-absolute-error vs. top-k% of the patches for error analyses between slides’ predictions and pathologists’ percent tumor nuclei estimates in (**g**) and slides’ predictions and genomic tumor purity values in (**h**).

In TCGA cohorts, pathologists’ percent tumor nuclei estimates were usually higher than genomic tumor purity values (Supp. 2.2). We investigated the probable causes of that difference. Our findings suggested that pathologists might have selected high tumor content regions to estimate percent tumor nuclei, which might have caused high percent tumor nuclei estimates (Figure 4g). We also observed that besides selecting the region-of-interest, its size is also crucial for some cancer types.

Our MIL models learned discriminant features for cancerous vs. normal histology while being trained on the weak labels of genomic tumor purity values. By conducting clustering over these features, we successfully obtained cancerous vs. normal segmentation maps for H&E stained slides of the TCGA LUAD cohort. Our qualitative validation showed that our segmentation is correct (Figure 1c and Figure 5).

**Figure 5:**
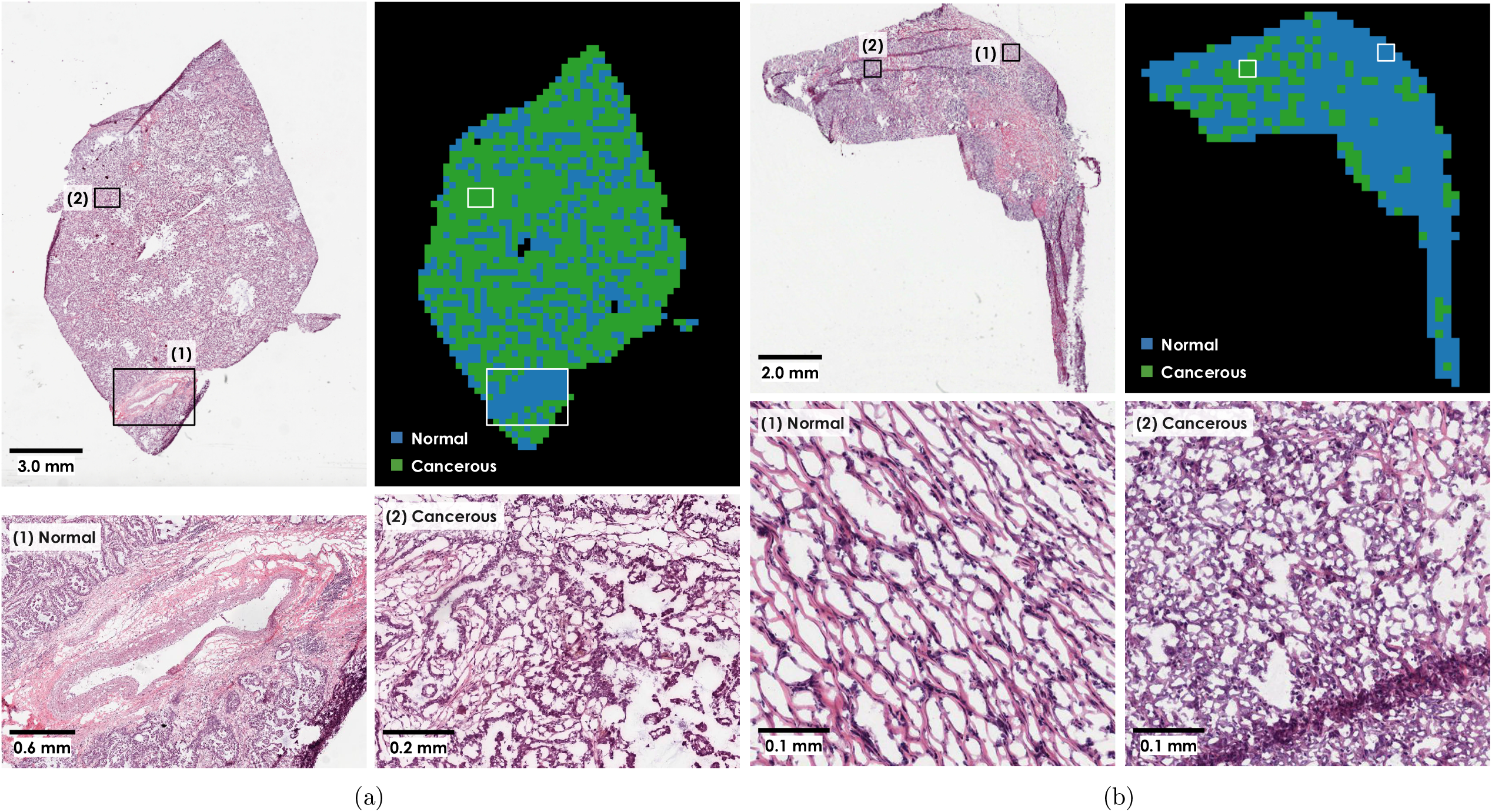
Cancerous vs. normal segmentation maps obtained by performing a clustering over the features extracted by the trained MIL model’s feature extractor module are consistent with LUAD histopathology. We show H&E stained WSIs, color-coded segmentation maps, and example zoom-in areas for two slides in the test set of LUAD cohort: (a) TCGA-73-4675-01A-01-TS1 and **(b)**TCGA-50-6590-01A-01-BS1.

Lastly, as an essential property of tumor purity predictors, we showed that our MIL models classified samples into tumor vs. normal almost perfectly in all cohorts (Figure 1d and Figure 3b).

## Results

We formulate predicting tumor purity of a sample from its H&E stained histopathology slides as a MIL task. The sample’s top and bottom slides are cropped into many patches, and these patches are collected to form a bag. Then, the task is to predict the bag level label of tumor purity. To achieve this task, we develop a novel MIL model consisting of three modules: *feature extractor* module, *MIL pooling filter*, and *bag-level representation transformation* module (Figure 1a). We use neural networks to implement the *feature extractor* module and the *bag-level representation transformation* module to parameterize the learning process fully (Methods). As the *MIL pooling filter*, we use our novel ‘distribution’ pooling filter. It is superior to standard pooling filters (like mean and max pooling) regarding the amount of information captured while obtaining bag-level representations [49]. Given a bag of patches, the *feature extractor* module extracts a feature vector for each patch inside the bag. Then, thanks to its superiority, the ‘distribution’ pooling filter obtains a strong bag-level representation by estimating the marginal distributions of the extracted features. Finally, the *bag-level representation transformation* module predicts tumor purity. This system of neural network modules is end-to-end trainable (see Methods for details).

In this study, there were ten different cohorts from TCGA and a local cohort from Singapore. Each TCGA cohort had more than 400 patients, and the Singapore cohort had 179 patients, such that each patient had both histopathology slides and corresponding genomic sequencing data (Table 1). In each cohort, we evaluated the performance of our trained MIL model on the data of completely unseen patients in the hold-out test set to simulate clinical workflow. In other words, each patient in the test set is like a new patient walking into the clinic in a real-world clinical setup [50].

### The MIL model’s tumor purity predictions correlate significantly with genomic tumor purity values

To evaluate our models’ performance in 10 different TCGA cohorts, correlation analyses between genomic tumor purity values (obtained from ABSOLUTE [18]) and our MIL models’ predictions are conducted. Spearman’s rank correlation coefficient is used as the performance metric.

In 8 cohorts, namely BRCA, GBM, LGG, LUAD, LUSC, OV, PRAD, and UCEC, we obtained significant correlations (P < 0.05) between genomic tumor purity values and our models’ predictions from digital histopathology slides (Figure 2). While the minimum Spearman’s *ρ_mil_* = 0.418 (P = 4.1e-05; 95% CI: 0.226 - 0.574) was obtained in the LGG cohort, the maximum Spearman’s *ρ_mil_* = 0.655 (P = 4.6e-24; 95% CI: 0.547 - 0.743) was obtained in the BRCA cohort.

We repeated the same analyses between genomic tumor purity values and pathologists’ percent tumor nuclei estimates (Supp. 2.1). While the minimum Spearman’s *ρ_path_* = 0.240 (P = 2.7e-02; 95% CI: 0.009 - 0.446) was obtained in the THCA cohort, the maximum Spearman’s *ρ_path_* = 0.344 (P = 9.8e-04; 95% CI: 0.139 - 0.531) was obtained in the UCEC cohort. There was no significant correlation in the GBM and LGG cohorts. Hence, the minimum correlation value obtained with MIL predictions (*ρ_mil_* = 0.418 in the LGG cohort) was higher than the maximum correlation value obtained with pathologists’ percent tumor nuclei estimates (*ρ_path_* = 0.344 in the UCEC cohort). This implies that MIL predictions are more consistent with genomic tumor purity values than the pathologists’ percent tumor nuclei estimates.

Moreover, we conducted statistical tests on correlation coefficients to compare: (i) our MIL models’ predictions and (ii) pathologists’ percent tumor nuclei estimates. We used the Fisher’s z transformation based method of Meng et al. [51]. We compared two methods only when there was a significant correlation for both methods in a cohort (Supp. Table 21). We observed that correlation coefficients obtained from MIL predictions were significantly better than ones obtained from pathologists’ estimates in all cohorts except LUSC and PRAD. For these cohorts, two methods performed on par (*P_comp_* = 1.7*e* – 01 > 0.05 for the LUSC and *P_comp_* = 2.0*e* – 01 >0.05 for the PRAD) in the test sets.

### MIL models’ predictions have lower mean absolute error than percent tumor nuclei estimates

Apart from Spearman’s correlation coefficients, we also checked the mean-absolute errors between genomic tumor purity values and MIL models’ predictions, and genomic tumor purity values and pathologists’ percent tumor nuclei estimates (see Supp. 2.2 for the complete analysis).

In the analyses of MIL predictions, the minimum and maximum mean-absolute-error values of *μ_e_mil__* = 0.105 (standard deviation *σ_e_mil__* = 0.091) and *μ_e_mil__* = 0.173 (*σ_e_mil__* = 0.154) were obtained in the OV cohort and the PRAD cohort, respectively. On the other hand, in the analyses of pathologists’ percent tumor nuclei estimates, the minimum and maximum mean-absolute-error values of *μ_e_path__* = 0.132 (*σ_e_path__* = 0.124) and *μ_e_path__* = 0.280 (*σ_e_path__* = 0.151) were obtained in the UCEC cohort and the LUAD cohort, respectively. In all cohorts, percent tumor nuclei estimates were generally higher than genomic tumor purity values.

Similar to our comparison in correlation analyses, we compared two methods based on absolute errors in the test sets of different cohorts. We used the Wilcoxon signed-rank test [52] on absolute error values for tumor samples in the test sets (Supp. Table 32). Absolute error values in MIL predictions were significantly lower than ones in pathologists’ percent tumor nuclei estimates in all cohorts except the LGG cohort. Two methods performed similar *(P_comp_* = 5.4*e* – 02 > 0.05) in the test set of the LGG cohort.

Our findings in correlation and absolute error analyses are summarized in Figure 3a. We observed that MIL predictions had lower mean-absolute-error and higher Spearman’s correlation coefficient than pathologists’ percent tumor nuclei estimates.

### The MIL model predicts tumor purity from H&E stained slides of FFPE sections in the Singapore cohort

Our MIL models successfully predicted tumor purity from H&E stained digital histopathology slides of fresh-frozen sections in different TCGA cohorts. Besides, we evaluated their performance on slides of formalin-fixed paraffin-embedded (ffpe) sections in a local Singapore cohort consisting of 179 lung adenocarcinoma patients. Similar to TCGA cohorts, we segregated data at the patient level (Table 1).

We used transfer learning and initialized the model with the weights of the MIL model trained on the TCGA LUAD cohort. Then, we freeze the weights of all layers in the network except the first convolutional layer in the feature extractor module (Figure 1a). This helped the network adapt the first layer weights to learn the tissue morphology in slides of ffpe sections, which were different from fresh-frozen sections (Supp. 3). Note that while the ffpe method preserves morphology better and is the routine in histopathology, the fresh-frozen method preserves nucleic acids better and is preferred for molecular analysis [53].

Similar to the performance in the TCGA LUAD cohort, we obtained a Spearman’s *ρ_mil_* = 0.554 (P = 4.6e-04; 95% CI: 0.283 - 0.745) and the mean-absolute-error of *μ_e_mil__* = 0.120 (*σ_e_mil__* = 0.091) in the test set of the Singapore LUAD cohort (Figure 2i, Figure 3a, and Supp. 3 for the complete analysis). There were substantial differences between the TCGA and Singapore LUAD cohorts, such as tissue preservation method (fresh-frozen vs. ffpe) and ancestry of patients (European vs. East Asian). However, our MIL model successfully predicted tumor purity from slides of ffpe sections using transfer learning with minimal training only in the first convolutional layer of the feature extractor module. The results suggested that our MIL models learned robust features for tumor purity prediction tasks at the higher levels of the network. We also checked the performance of the TCGA LUAD model directly on the Singapore LUAD cohort used as an external validation set (Supp. 3). Nevertheless, we could not get as good results as we got in transfer learning, which highlighted the necessity of adapting the weights of the first layer in feature extractor to ffpe slides.

### Tumor purity varies spatially within a sample: top and bottom slides of a sample are different in tumor purity

Intra-tumor heterogeneity is a well-known phenomenon in solid cancers [54–58]. It results in therapeutic failure and drug resistance [59]. We checked whether it is observable from tumor purity predictions of the trained MIL model on the top and bottom slides of a sample (see Methods). For each slide of a tumor sample with both top and bottom slides in a cohort, 100 bags are created by randomly sampling from available patches of the slide and predictions are obtained from the trained MIL model. Then, the predictions of two slides are statistically compared using the Wilcoxon signed-rank test [52] (see Supp. 2.3 for the complete analysis).

Figure 3c shows the box plot of p-values obtained from the statistical tests in each cohort’s test set. There is a significant difference between the MIL predictions on the top and bottom slides of the same tumor sample. In all cohorts, at least 75% of samples have p-value *P* < 1.0*e* – 08 and at least 95% of samples have p-value P < 0.05. Hence, we conclude that there is a variation in tumor purity between the top and bottom sections of a tumor sample, i.e., tumor purity varies spatially within the sample.

The degree of spatial variation in tumor purity is different for different cancer types (Supp. 2.3). The UCEC, LGG, and GBM cohorts had the lowest mean absolute differences (*μ_d_abs__*) between top and bottom slides’ predictions (*μ_d_abs__* ≤ 0.090), i.e., they were the most spatially homogeneous cancers among all cohorts (Supp. Table 41). On the other hand, the PRAD cohort had the highest mean absolute difference (*μ_d_abs__* = 0.144), i.e., it was the most spatially heterogeneous cancer in tumor purity.

### Predicting the tumor purity of a sample by using both top and bottom slides is better than using only one slide

We checked if there is a significant difference between predicting a sample’s tumor purity by using both slides (top and bottom) and using only one slide. For a tumor sample with two slides in a cohort, let *p_smpl_* be genomic tumor purity value of the sample; 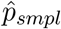 be tumor purity prediction obtained from trained MIL model by using both of the slides together; 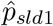 and 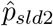 be tumor purity predictions obtained from trained MIL model for individual slides. We compared the absolute error of sample level prediction 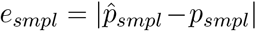 and the expected value of absolute errors of slide level predictions 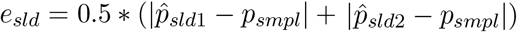 (see Supp. 2.4 for complete analysis). We used the Wilcoxon signed-rank test [52] on the difference of *e_smpl_* – *e_sld_* (Supp. Table 50). Note that the PRAD (n=21) and UCEC (n=23) cohorts were excluded from this study due to few samples with two slides.

In the test sets of BRCA, LUAD, LUSC, and OV cohorts, using both slides for tumor purity prediction gave better results in terms of absolute error (Figure 3d). However, in the test sets of GBM and LGG cohorts, there was no significant difference using both slides or one slide alone. Indeed, this is not surprising since they had the lowest mean absolute differences between the slides’ predictions (Supp. Table 41), i.e., the most spatially homogeneous tumors. In fact, when both slides are the same, sample level prediction and slide predictions would be the same.

We conclude that predicting a sample’s tumor purity using both the top and bottom slides together is better than using only one of them whenever possible.

### Spatial tumor purity map analysis reveals the probable cause of pathologists’ high percent tumor nuclei estimates

Pathologists’ percent tumor nuclei estimates were generally higher than genomic tumor purity values for all TCGA cohorts in our analysis (Supp. 2.2). We hypothesized that inappropriate size and selection of region-of-interest might be the cause. We obtained tumor purity maps by our trained MIL models in different TCGA cohorts and conducted error analysis over them to test our hypothesis.

We followed the same procedure in Smits et al. [5] to simulate pathologists’ percent tumor nuclei estimation. Tumor purity is predicted over a region-of-interest of 1*mm* × 1*mm* around each patch in a slide, which corresponds to 16 patches at 20× zoom level (each patch is around 256*μm* × 256*μm* at the specimen level) (Figure 4a). Then, the predicted value is assigned to the patch in the tumor purity map (Figure 4b). We also obtained tumor purity maps for slides in the Singapore cohort (Figure 4d and Figure 4e).

We observed that a tumor purity map shows variation within the slide, which implies that region-of-interest selection is crucial in pathologists’ percent tumor nuclei estimation. Since tumor purity was higher in pathologists’ percent tumor nuclei estimates, we investigated whether pathologists might have selected high tumor content regions over the slides for percent tumor nuclei estimation. The highest prediction in a slide’s tumor purity map was used as the slide’s tumor purity value. Then, error analyses were conducted over the slides’ tumor purity values compared to pathologists’ percent tumor nuclei estimates and genomic tumor purity values. The error analyses were repeated by gradually extending the region-of-interest such that a slide’s tumor purity was calculated as the average of top-k% of the patches with the highest scores (*k* = 0, …, 100) in the slide’s tumor purity map.

We discovered that the mean-absolute-error between the slides’ predictions and pathologists’ percent tumor nuclei estimates increases as we extend the region-of-interest to cover the lower tumor purity regions (Figure 4g). These observations suggested that pathologists may tend to select high tumor content regions to estimate percent tumor nuclei. The LGG and UCEC cohorts may look exceptional with almost constant mean-absolute-error plots. However, this is expected since these two cohorts’ samples have high genomic tumor purity values (Supp. 1.2), so the variation within the slides is very low. The PRAD cohort’s plot also has a different pattern than the others. It has an initial decrease and an increase in the latter stages, emphasizing the importance of the region-of-interest size. The pathologists may need to analyze a bigger region-of-interest depending on the morphology of the tissue origin to reach a certain nuclei count while estimating percent tumor nuclei. The PRAD may be one of them due to the glandular structure of the prostate.

Furthermore, as the region-of-interest grows, the mean-absolute-error between the slides’ predictions and genomic tumor purity values decreases (Figure 4h). Indeed, this is expected since our MIL models converge to their original performance of prediction over the whole slide (Figure 2). It is even more evident in the LUSC and OV cohorts. The error decreases initially but increases later since our MIL models underestimated the tumor purity compared to genomic tumor purity values in these cohorts.

### The MIL model learns discriminant features for cancerous vs. normal tissue histology

We explored the capability of our MIL model’s feature extractor on learning discriminant features for cancerous vs. normal tissue histology while being trained on sample-level genomic tumor purity labels. For each patient having both tumor and matching normal samples, features of patches cropped over the slides of the tumor and normal samples were extracted using the trained feature extractor module of the MIL model. Then, slide-level cancerous vs. normal segmentation maps were obtained by performing a clustering over the extracted feature vectors (see Methods). The resolution of segmentation was at the patch level, and each patch was around 256*μm* × 256*μm* at the specimen level.

In the test set of the LUAD cohort, there were 33 patients both with tumor and matching normal samples. We constructed slide-level segmentation maps for these patients (Figure 5). We observed that segmentation maps were consistent with the LUAD histopathology during the qualitative assessment of the segmentation maps. While healthy tissue components, like blood vessels, stroma regions, and normal tissue structures, were labeled normal, regions invaded by neoplastic cells were labeled cancerous. Hence, we qualitatively validated that our MIL model learned discriminant features for cancerous vs. normal tissue histology in LUAD from sample-level genomic tumor purity labels without requiring exhaustive annotations from pathologists.

### The MIL model successfully classifies samples into tumor vs. normal

Tumor vs. normal discrimination is one of the essential properties of a tumor purity predictor. To be able to predict tumor purity correctly, it must learn to discriminate between tumor and normal. We checked our MIL model’s performance in the tumor vs. normal sample classification task.

Tumor purity predictions for all samples in the test set of each cohort were obtained and a receiver operating characteristic (ROC) curve analysis was conducted. Then, the area under the ROC curve (AUC) was calculated and a 95% confidence interval was constructed using the percentile bootstrap method [60]. Note that GBM and LGG cohorts were excluded from analysis since there were no normal slides in these cohorts.

Our MIL models successfully discriminated tumor samples from normal samples in all cohorts with AUC values greater than or equal to 0.927 (Figure 3b). We got the minimum and maximum AUC values of 0.927 (95% CI: 0.826 - 0.993) and 1.000 (95% CI: 1.000 - 1.000) on the test sets of THCA and BRCA cohorts, respectively. Note that although we did not get a strong correlation between genomic tumor purity values and MIL predictions in the test sets of KIRC and THCA cohorts, our models successfully classified samples into tumor vs. normal in these cohorts.

Furthermore, we obtained an AUC value of 0.991 (95% CI: 0.975 - 1.000) on the test set of the LUAD cohort. Our model outperformed the classical image processing and machine learning-based method of Yu et al. (AUC: 0.85) [61] and the DNA plasma-based method of Sozzi et al. (AUC: 0.94) [62]. Besides, our model performed on par with the deep learning model of Coudray et al. [63] (AUC: 0.993), which was trained on tumor vs. normal classification, and the deep learning model of Fu et al. [64] (AUC: 0.977 with 95% CI: 0.976 - 0.978), which was fine-tuned on pathologists’ percent tumor nuclei estimates in a transfer learning setup. However, there is one concern about the dataset preparation methods of Coudray et al. [63] and Fu et al. [64]. They obtained the datasets by segregating data either at slide level [63] or at patch level [64]. These data segregation methods might lead to a severe data leakage problem, and the models’ performance might be illusory.

## Discussion

Tumor purity is a crucial prognostic biomarker. It also affects the quality of molecular data acquisition and analysis. However, percent tumor nuclei estimation by pathologists is tedious and time-consuming. Besides, pathologists’ estimates suffer from inter-observer variability. To overcome these challenges, we developed a novel MIL model with a distribution pooling filter. It predicted tumor purity from H&E stained histopathology slides of fresh-frozen and formalin-fixed paraffin-embedded sections in different TCGA cohorts and a Singapore cohort, respectively. The predictions were consistent with genomic tumor purity values, and they outperformed pathologists’ percent tumor nuclei estimates in the TCGA cohorts.

Hence, our MIL models can be utilized in sample selection for molecular analysis, which will help reduce pathologists’ workload and decrease inter-observer variability. Moreover, spatially resolved tumor purity maps obtained using our MIL models can substantially contribute to a better understanding of the tumor microenvironment. Lastly, our models’ predictions can be used as prognostic biomarkers to stratify patients.

### Weak tumor purity labels innately necessitated a MIL approach

The necessity of automated tumor purity prediction from H&E stained digital histopathology images has recently been recognized. There have been few studies in different cancer types, like breast cancer [41–44], lung cancer [40, 44], and colon cancer [39, 44]. They were all patch-based methods, and they relied on expensive pixel-level annotations. Some of them extracted intensity and morphology-based hand-crafted features over the annotated regions and employed traditional machine learning methods, such as support vector machines (SVMs) [40, 41] or random forest [44], to classify the patches into malignant vs. benign. Then, they estimated tumor purity. Besides, recent two studies used deep neural networks (DNNs) for tumor cellularity estimation in image patches [42, 43]. One extracted the patch features using a pre-trained DNN and trained decision trees and SVMs to predict the tumor cellularity [42]. The other one predicted tumor cellularity directly from image patches [43].

The limiting factor in these studies was the requirement for pixel-level annotations, which usually do not exist since it is a time-consuming and tedious process. However, sample-level weak labels are easier to obtain from pathology reports, electronic health records, or different data modalities. This study used genomic tumor purity values obtained from genomic sequencing data by ABSOLUTE [18] as sample-level weak labels.

While previous studies worked on few cancer types with relatively few patients (like 10 patients [41] or 64 patients [42, 43]), this study conducted a pan-cancer study on 10 different TCGA cohorts, where each cohort had more than 400 patients. In this study, genomic tumor purity values as ‘accurate’ labels helped us obtain strong models. However, unlike pixel-level annotations providing whether each cell is cancerous or normal, the genomic tumor purity of a sample tells us only the proportion of cancer cells within the sample, so it is a sample-level weak label. Besides, a sample is best represented using both top and bottom slides. Therefore, training a machine learning model, which predicts sample-level tumor purity, using weak genomic tumor purity labels innately necessitated a MIL approach. This approach represented a sample as a bag of patches from the sample’s slides and used the sample’s genomic tumor purity value as the bag’s label.

### The sources of error in MIL predictions

Our MIL models successfully predicted tumor purity (Figure 2). However, they slightly deviated from the genomic tumor purity values. There may be different sources of prediction errors. While some of them can be eliminated, some are inevitable.

Firstly, we have fewer patients in our data sets than traditional deep learning data sets containing millions of independent samples [65]. There are around 500 patients per dataset on average, so around 300 patients per training set. Considering the complexity of cancer, our MIL models effectively captured features that distinguish cancerous vs. normal. We also expect that the performance will improve with the increasing number of patients. Indeed, we obtained the best performance in our largest cohort of BRCA, which had 559 patients in the training set.

Secondly, our MIL model uses histopathology slides from the top and bottom sections of the tumor portion. We have already shown the variation in tumor purity between the top and bottom sections of the tumor samples. Thus, for samples with only one slide, the prediction error is expected to be higher.

Lastly, our model’s predictions are based on morphology in H&E stained histopathology slides. However, genomic tumor purity values were based on DNA data, and all the effects of genetic changes (so, the genomic tumor purity changes) may not be observable from the slides due to the selective dying characteristics of H&E staining. Besides, it is a well-known fact that DNA manifests morphology, which is the basis of this study as well. Nevertheless, there is a long way from DNA to morphology. Some factors, like epigenetics, cell differentiation, or gene expression stochasticity, may hinder the effects of some mutations occurring in DNA from manifesting themselves in morphology [66].

### Superiority of MIL predictions over percent tumor nuclei estimates

Comparing with the percent tumor nuclei estimates by pathologists, our MIL models’ predictions gave a higher correlation and lower mean-absolute-error with genomic tumor purity values (Figure 3a). One of the primary reasons for this superiority is that the MIL models were trained directly on genomic tumor purity values, which enabled the MIL models to learn associated features.

Another reason might be that pathologists concentrate more on tumor cells than infiltrating normal cells within the tumor, which may result in missed normal tissue components. Moreover, cancer cells are usually enlarged. They occupy more space than normal tissue components, stromal cells, and infiltrating lymphocytes, which may create an implication of high tumor content [17]. Pathologists may fail to incorporate this effect in their estimates correctly and may overestimate percent tumor nuclei. Indeed, this was the case in the cohorts we analyzed (Supp. 2.1).

Finally, while our MIL models predict tumor purity over the whole slide, pathologists estimate the percent tumor nuclei by analyzing some selected region-of-interest over the slide. Therefore, the size and selection of the region-of-interest might cause the overestimation in pathologists’ percent tumor nuclei estimates (Figure 4g).

### Spatially resolved tumor purity maps can complement spatial-omics

We obtained tumor purity maps showing the variation of tumor purity in slides using our trained MIL models (Figure 4b and Figure 4e). They can potentially help understand the interaction of cancer cells with other tissue components (like normal epithelial, stromal, and immune cells) in the tumor microenvironment, which is a key player in tumor formation and primary determinant of therapeutic response [38, 67]. Furthermore, they can complement spatial-omics analyses revealing the spatial distribution of omics features [68–70].

### The MIL model learns discriminant features from weak labels

Our MIL model in the TCGA LUAD cohort learned discriminant features that can classify cancerous and normal tissues correctly in LUAD dataset (Figure 5). Note that our MIL model learned discriminant features from genomic tumor purity values (which are sample-level weak labels) without requiring pixel-level annotations from pathologists, which are expensive and tedious. It is pretty promising to explore weak labels further for new biomarkers in cancer studies. In other words, the question of ‘How strong are the weak labels?’ is still a valid research question to be explored in digital histopathology.

### Limitations

Our MIL models, by design, apply to any tumor sample with H&E stained histopathology slides. We tested them on tumor samples with a broad range of tumor purity values. However, checking their performance on samples with low tumor content (where ABSOLUTE cannot determine the tumor purity values accurately) would strengthen the applicability of our MIL models. It is reserved for future work.

We evaluated our MIL models on hold-out test sets to simulate real-world clinical workflow and obtained successful results. Besides, our analysis on the Singapore LUAD cohort using transfer learning with minimal training for domain adaptation showed that our MIL models learned robust features for tumor purity prediction tasks. However, we could not validate our models on external cohorts due to differences between fresh-frozen and formalin-fixed paraffin-embedded tissue preservation methods, which might further consolidate their robustness.

Lastly, our MIL models are deep learning based, and deep learning algorithms perform better with more data. Training of the models with larger cohorts would help to improve the model performance by better capturing patient-to-patient variations.

## Methods

### Data Sets

We downloaded H&E stained fresh-frozen section histopathology slides and corresponding genomic sequencing data for ten different cohorts in TCGA (BRCA, GBM, KIRC, LGG, LUAD, LUSC, OV, PRAD, THCA, and UCEC). We selected these cohorts since they have more than 400 patients with both histopathology slides and corresponding genomic sequencing data in TCGA (Table 1). Each patient had a tumor sample, and some patients also had matching normal samples.

In TCGA, each sample was chopped into portions following the TCGA Standard Operating Procedures [71–74]. Then, one of the portions was sequenced for genomic analysis. One or two associated histopathology slides (namely top and bottom slides) were also prepared from the top and bottom sections of the same portion.

We also collected digital histopathology slides of an East Asian cohort consisting of 179 lung adenocarcinoma patients in Singapore. The genomic data for this cohort is publicly available from OncoSG (https://src.gisapps.org/OncoSG/) under dataset ‘Lung Adenocarcinoma (GIS, 2019)’. Although slides are not publicly available, this site serves representative histological images for each patient. In the Singapore cohort, only one slide was prepared for each tumor sample from the top section of the tissue used for sequencing, and there were no normal samples.

In each cohort, we randomly segregated the data at the patient level (i.e., slides from the same patient should be in the same set) into training, validation, and test sets, which had similar tumor purity distributions (Supp. 1.2). Note that segregating data at the patient level is crucial to prevent data leakage while training machine learning models [50]. The training set was used to train the machine learning model, the validation set was used to choose the best model, and the test set was held out as unseen data for evaluation of the best model. The list of patients and slides in each set are given in Supp. File 1.

Tissue regions inside histopathology slides were detected by applying OTSU thresholding, image dilation, median filtering, and hole-filling, respectively. Over the detected tissue regions, non-overlapping 512 × 512 RGB images at 20× zoom level (specimen-level pixel size, 0.5*μm* × 0.5*μm*) were cropped. For each cohort, detailed information about the data in training, validation, and test sets are given in Supp. 1.

### MIL Model

#### Problem formulation and notation

The objective is to predict a bag label *Y* for a given bag of instances *X* = {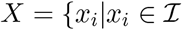, *i* = 1, 2, …, *N*} where 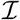 is the instance space and *N* is the number of instances inside the bag. Here, a bag label *Y* is the genomic tumor purity of a sample, and a bag *X* is a collection of cropped patches over the sample’s slides.

Let 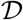 be a MIL dataset such that 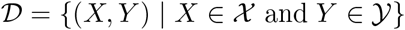, where 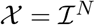 is the bag space and 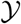 is the bag label space. Given any pair 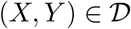, our objective is to predict bag label *Y* for a given bag of instances *X* = {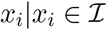, *i* = 1, 2, …, *N*}. Let 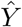 be the predicted bag label of *X*. To obtain 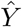, we designed a novel MIL framework consisting of three stages.

The first stage is a *feature extractor* module 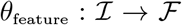, where 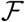 is the feature space. For each *x_i_* ∈ *X*, it takes *x_i_* as input, extracts *J* features and outputs a feature vector: 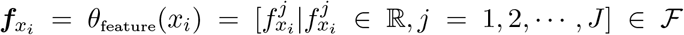 where 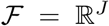. Let 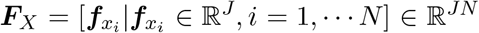 be a feature matrix constructed from extracted feature vectors such that *i^th^* column corresponds to ***f**_x_i__*.

The second stage is a *MIL pooling filter* module 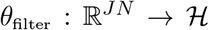, where 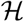 is the bag-level representation space. It takes the feature matrix ***F**_X_* as input and aggregates the extracted feature vectors into a bag-level representation: 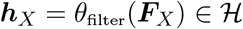.

The last stage is a *bag-level representation transformation* module 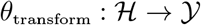. It transforms the bag level representation into the predicted bag label: 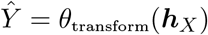.

We use neural networks to implement *θ*_feature_ and *θ*_transform_ so that we can fully parameterize the learning process. For *θ*_filter_, we use our novel ‘distribution’ pooling filter [49], which we have shown to be superior to point-estimate based MIL pooling filters, like max-pooling or mean-pooling. This system of neural networks is end-to-end trainable.

#### Distribution Pooling Filter

We have defined the family of distribution-based pooling filters in [49] as: Given a feature matrix ***F**_X_* = [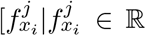, *i* = 1, 2, …, *N* and *j* = 1, 2, …, *J*] obtained from a bag *X* = {*x*_1_, *x*_2_, …, *x_N_*}, its bag level representation is obtained by estimating a marginal distribution over each extracted feature. Let 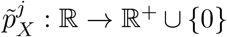 be the estimated marginal distribution obtained over *j^th^* extracted feature and 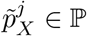 where 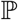 is the set of all possible marginal distributions. 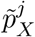 is calculated by using kernel density estimation [75], which employs a Gaussian kernel with standard deviation *σ*, as shown in the Eq. 1. Each instance has two attention based weights, feature weight *α_i_* and kernel weight *β_i_*, obtained from neural network modules. Hence, the bag level representation 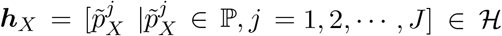 where 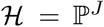. Note that the estimated marginal distributions are uniformly binned during training neural network models for computational purposes.

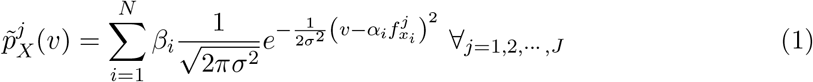

In [49], we formally proved that the distribution-based pooling filters are superior to the point estimate-based counterparts (like max and mean pooling) regarding the amount of information captured while obtaining bag-level representations. Then, we empirically showed that models with distribution-based pooling filters perform equal or better than that with point estimate-based pooling filters on distinct real-world MIL tasks.

In this study, we used standard deviation of *σ* = 0.05 and the estimated marginal distributions were uniformly binned into 21 bins. Note that attention weights in ‘distribution’ pooling were fixed to *α_i_* = 1 ∀_*i*_ and *β_i_* = 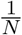 ∀_*i*_ where *N* is the number of instances per bag.

#### Neural network architectures and hyper-parameters

We used a ResNet18 [76] model as the *feature extractor* module and a three-layer multi-layer-perceptron as the *bag-level representation transformation* module.

During the training of the models, we prepared bags on the go. A bag was created by randomly sampling 200 patches (instances) from all available patches previously cropped over a sample’s slides. The patch size was 512 × 512. Data augmentation (random cropping with a size of 299 × 299 and random horizontal/vertical flipping) was also applied on the patches. We extracted 128 features for each instance inside the bag.

### Training of MIL Model

We trained the MIL model on samples of patients in the training set. We used a bag of patches from slides of the tumor sample as the input and tumor purity value obtained from genomic sequencing data by ABSOLUTE [18] as the ground-truth label (Supp. 1.2). Genomic tumor purity values were extracted from publicly available data in the Genomic Data Commons (https://gdc.cancer.gov/about-data/publications/pancanatlas) under filename “TCGA_mastercalls.abs_tables_JSedit.fixed.txt” for the TCGA cohorts and in the OncoSG (https://src.gisapps.org/OncoSG/) under dataset ‘Lung Adenocarcinoma (GIS, 2019)’ for the Singapore cohort. We also used the matching normal sample of the same patient whenever available to enable our model to capture the information related to normal tissue histology. We assigned a tumor purity value of 0.0 to a matching normal sample as the ground-truth label. Note that there were no normal samples in the Singapore cohort.

We initialized the neural networks randomly and trained them end-to-end. We trained models using the ADAM optimizer with a learning rate of *lr* = 0.0001 and *L*2 regularization on the weights with a weight decay of *weight_decay* = 0.0005. The batch size was 1. We used absolute error as the loss function and employed early-stopping based on loss in the validation set to avoid overfitting. Then, we evaluated our model on the unseen test set.

### Predicting Tumor Purity of a Sample

We created 100 bags for each sample in the test set and obtained tumor purity predictions from the trained model. We used the average of 100 predictions as the sample’s tumor purity prediction during performance evaluation.

### Segmentation of Histopathology Slides

Understanding the structure of tumor composition is essential. We developed a cancerous vs. normal segmentation algorithm in the TCGA LUAD cohort to visually inspect the structure of the tumor microenvironment.

In principle, a tumor purity predictor must learn discriminant features for cancerous and normal components in the tissue to predict tumor purity correctly. For each patient with a matching normal sample, we used the trained feature extractor module of our MIL model to extract features of patches cropped over the slides of the tumor and normal samples of the patient. Then, we clustered the patches using hierarchical clustering over the extracted feature vectors (Figure 1c). We calculated patient-specific distance thresholds in hierarchical clustering to capture inter-patient variations. Finally, we assigned a cancerous or normal label to each cluster based on the slide type (tumor slide or normal slide) that the patches within the cluster belong to (Supp. 4).

### Statistical Analysis

We obtained 95% confidence intervals for Spearman’s rank correlation coefficients and area under the receiver operating characteristic curves using the percentile bootstrap method [60].

To compare the performance of two methods (our MIL models’ predictions and pathologists’ percent tumor nuclei estimates), we used Fisher’s z transformation based method of Meng et al. [51] on Spearman’s rank correlation coefficients and Wilcoxon signed-rank test [52] on absolute error values.

All statistical tests were two-sided and statistical significance was considered when P < 0.05. We used scipy.stats (v1.4.1) python library for statistical tests.

## Data Availability

All TCGA datasets are publicly available. Manifest files can be obtained from the GitHub repository (https://github.com/onermustafaumit/SRTPMs) to download H&E stained digital histopathology slides using GDC Data Transfer Tool. Genomic tumor purity values can be downloaded from https://gdc.cancer.gov/about-data/publications/pancanatlas under filename “TCGA_mastercalls.abs_tables_JSedit.fixed.txt”. For the Singapore cohort, genomic tumor purity values and representative histological images are publicly available from OncoSG (https://src.gisapps.org/OncoSG/) under dataset ‘Lung Adenocarcinoma (GIS, 2019)’.

## Code Availability

The source code is available at https://github.com/onermustafaumit/SRTPMs. The repository provides a detailed step-by-step explanation, from downloading H&E stained digital histopathology slides to obtaining **S**patially **R**esolved **T**umor **P**urity **M**ap**s** (SRTPMs).

## Acknowledgements

This work is partly supported by the Biomedical Research Council of the Agency for Science, Technology, and Research, Singapore and the National University of Singapore, Singapore. We thank Valerie Yang for constructive comments on the manuscript.

## Author Contributions

M.U.O., W.K.S., and H.K.L. conducted machine learning study. J.C., E.R., A.N.K., J.J.S.A., W.Z. and A.J.S. performed genomic analysis. D.S.W.T. designed the clinical study of Singapore cohort. A.J., S.Y.H., A.T., X.M.C. and T.K.H.L. conducted histopathological work and prepared samples and slides in the Singapore cohort.

