## Supplementary Material for "Obtaining Spatially Resolved Tumor Purity Maps Using Deep Multiple Instance Learning In A Pan-cancer Study"

### 1 Summary of Datasets in TCGA Cohorts

#### 1.1 The number of samples, slides, and patches in datasets

The number of samples, slides, and patches in each set are given for: BRCA cohort in Table 1, GBM cohort in Table 2, KIRC cohort in Table 3, LGG cohort in Table 4, LUAD cohort in Table 5, LUSC cohort in Table 6, OV cohort in Table 7, PRAD cohort in Table 8, THCA cohort in Table 9, UCEC cohort in Table 10.

Each patient has only one tumor sample and one normal sample if available. Note that "tumor slide" and "normal slides" refer to the slides of tumor samples and normal samples, respectively. Similarly, "tumor patches" and "normal patches" refer to patches cropped over "tumor slides" and "normal slides", respectively.

Table 1: **BRCA cohort.** The number of samples, slides, and patches in datasets.

| | # samples | | | # slides | | | # patches | | |
|---|---|---|---|---|---|---|---|---|---|
| dataset | normal | tumor | total | normal | tumor | total | normal | tumor | total |
| training | 76 | 559 | 635 | 177 | 763 | 940 | 41,870 | 419,954 | 461,824 |
| validation | 27 | 185 | 212 | 68 | 259 | 327 | 24,553 | 150,481 | 175,034 |
| test | 30 | 185 | 215 | 67 | 258 | 325 | 17,773 | 140,011 | 157,784 |
| all | 133 | 929 | 1,062 | 312 | 1,280 | 1,592 | 84,196 | 710,446 | 794,642 |

Table 2: **GBM cohort.** The number of samples, slides, and patches in datasets.

| | # samples | | | # slides | | | # patches | | |
|---|---|---|---|---|---|---|---|---|---|
| dataset | normal | tumor | total | normal | tumor | total | normal | tumor | total |
| training | 0 | 285 | 285 | 0 | 555 | 555 | 0 | 360,662 | 360,662 |
| validation | 0 | 95 | 95 | 0 | 178 | 178 | 0 | 124,979 | 124,979 |
| test | 0 | 94 | 94 | 0 | 184 | 184 | 0 | 133,008 | 133,008 |
| all | 0 | 474 | 474 | 0 | 917 | 917 | 0 | 618,649 | 618,649 |

Table 3: **KIRC cohort.** The number of samples, slides, and patches in datasets.

| | # samples | | | # slides | | | # patches | | |
|---|---|---|---|---|---|---|---|---|---|
| dataset | normal | tumor | total | normal | tumor | total | normal | tumor | total |
| training | 220 | 261 | 481 | 276 | 501 | 777 | 285,947 | 393,683 | 679,630 |
| validation | 71 | 85 | 156 | 84 | 165 | 249 | 90,094 | 122,039 | 212,133 |
| test | 73 | 89 | 162 | 94 | 175 | 269 | 90,842 | 139,903 | 230,745 |
| all | 364 | 435 | 799 | 454 | 841 | 1295 | 466,883 | 655,625 | 1,122,508 |

Table 4: **LGG cohort.** The number of samples, slides, and patches in datasets.

| | # samples | | | # slides | | | # patches | | |
|---|---|---|---|---|---|---|---|---|---|
| dataset | normal | tumor | total | normal | tumor | total | normal | tumor | total |
| training | 0 | 273 | 273 | 0 | 380 | 380 | 0 | 210,371 | 210,371 |
| validation | 0 | 91 | 91 | 0 | 124 | 124 | 0 | 75,533 | 75,533 |
| test | 0 | 90 | 90 | 0 | 121 | 121 | 0 | 61,161 | 61,161 |
| all | 0 | 454 | 454 | 0 | 625 | 625 | 0 | 347,065 | 347,065 |

Table 5: **LUAD cohort.** The number of samples, slides, and patches in datasets.

| | # samples | | | # slides | | | # patches | | |
|---|---|---|---|---|---|---|---|---|---|
| dataset | normal | tumor | total | normal | tumor | total | normal | tumor | total |
| training | 101 | 266 | 367 | 120 | 418 | 538 | 67,174 | 296,382 | 363,556 |
| validation | 37 | 90 | 127 | 43 | 142 | 185 | 22,768 | 92,958 | 115,726 |
| test | 33 | 90 | 123 | 37 | 134 | 171 | 18,934 | 101,061 | 119,995 |
| all | 171 | 446 | 617 | 200 | 694 | 894 | 108,876 | 490,401 | 599,277 |

Table 6: **LUSC cohort.** The number of samples, slides, and patches in datasets.

| | # samples | | | # slides | | | # patches | | |
|---|---|---|---|---|---|---|---|---|---|
| dataset | normal | tumor | total | normal | tumor | total | normal | tumor | total |
| training | 132 | 273 | 405 | 198 | 432 | 630 | 104,170 | 339,703 | 443,873 |
| validation | 41 | 90 | 131 | 62 | 140 | 202 | 32,474 | 105,556 | 138,030 |
| test | 47 | 90 | 137 | 73 | 142 | 215 | 29,537 | 99,519 | 129,056 |
| all | 220 | 453 | 673 | 333 | 714 | 1,047 | 166,181 | 544,778 | 710,959 |

Table 7: **OV cohort.** The number of samples, slides, and patches in datasets.

| | # samples | | | # slides | | | # patches | | |
|---|---|---|---|---|---|---|---|---|---|
| dataset | normal | tumor | total | normal | tumor | total | normal | tumor | total |
| training | 53 | 310 | 363 | 92 | 620 | 712 | 39,215 | 667,145 | 706,360 |
| validation | 13 | 103 | 116 | 23 | 206 | 229 | 19,446 | 234,701 | 254,147 |
| test | 18 | 103 | 121 | 27 | 205 | 232 | 13,724 | 220,774 | 234,498 |
| all | 84 | 516 | 600 | 142 | 1,031 | 1,173 | 72,385 | 1,122,620 | 1,195,005 |

Table 8: **PRAD cohort.** The number of samples, slides, and patches in datasets.

| | # samples | | | # slides | | | # patches | | |
|---|---|---|---|---|---|---|---|---|---|
| dataset | normal | tumor | total | normal | tumor | total | normal | tumor | total |
| training | 72 | 258 | 330 | 72 | 324 | 396 | 49,001 | 205,163 | 254,164 |
| validation | 15 | 85 | 100 | 15 | 105 | 120 | 11,026 | 64,559 | 75,585 |
| test | 24 | 85 | 109 | 24 | 106 | 130 | 15,771 | 68,398 | 84,169 |
| all | 111 | 428 | 539 | 111 | 535 | 646 | 75,798 | 338,120 | 413,918 |

Table 9: **THCA cohort.** The number of samples, slides, and patches in datasets.

| | # samples | | | # slides | | | # patches | | |
|---|---|---|---|---|---|---|---|---|---|
| dataset | normal | tumor | total | normal | tumor | total | normal | tumor | total |
| training | 48 | 258 | 306 | 48 | 270 | 318 | 18,678 | 122,883 | 141,561 |
| validation | 18 | 85 | 103 | 18 | 87 | 105 | 6,395 | 37,772 | 44,167 |
| test | 17 | 85 | 102 | 17 | 86 | 103 | 5,161 | 38,620 | 43,781 |
| all | 83 | 428 | 511 | 83 | 443 | 526 | 30,234 | 199,275 | 229,509 |

Table 10: **UCEC cohort.** The number of samples, slides, and patches in datasets.

| | # samples | | | # slides | | | # patches | | |
|---|---|---|---|---|---|---|---|---|---|
| dataset | normal | tumor | total | normal | tumor | total | normal | tumor | total |
| training | 18 | 270 | 288 | 18 | 358 | 376 | 8,928 | 192,766 | 201,694 |
| validation | 4 | 90 | 94 | 4 | 119 | 123 | 1,004 | 65,596 | 66,600 |
| test | 10 | 89 | 99 | 12 | 112 | 124 | 7,427 | 56,262 | 63,689 |
| all | 32 | 449 | 481 | 34 | 589 | 623 | 17,359 | 314,624 | 331,983 |

##### 1.2 Genomic tumor purity values

We used genomic tumor purity values obtained using ABSOLUTE [1] as labels. Histogram plots of genomic tumor purity values of tumor samples in each set are given for: BRCA cohort in Figure 1, GBM cohort in Figure 2, KIRC cohort in Figure 3, LGG cohort in Figure 4, LUAD cohort in Figure 5, LUSC cohort in Figure 6, OV cohort in Figure 7, PRAD cohort in Figure 8, THCA cohort in Figure 9, UCEC cohort in Figure 10.

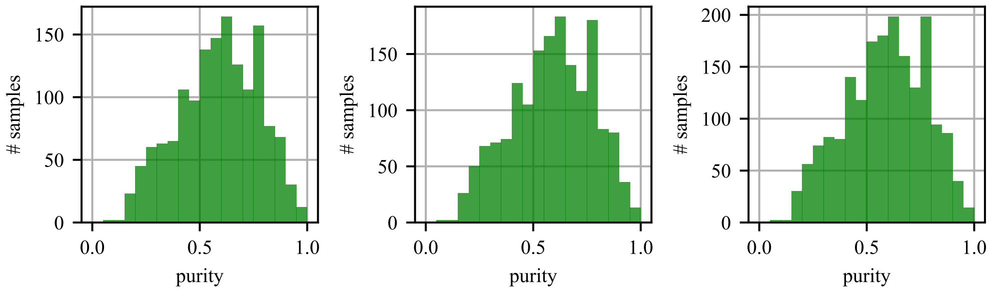


(a) training (b) validation (c) test

Figure 1: **BRCA cohort:** Genomic tumor purity histograms for training, validation, and test sets.

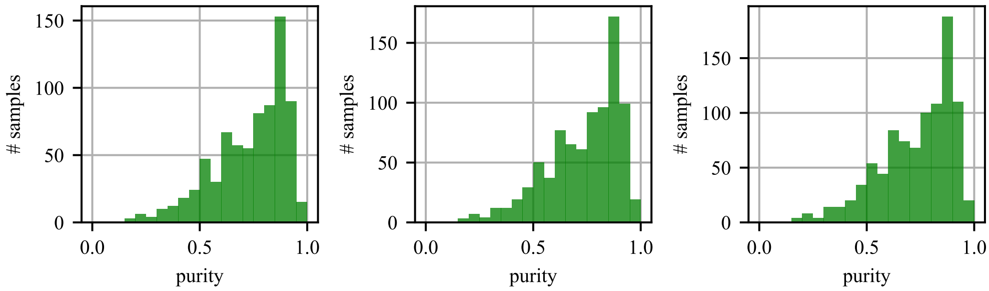


(a) training (b) validation (c) test

Figure 2: **GBM cohort:** Genomic tumor purity histograms for training, validation, and test sets.

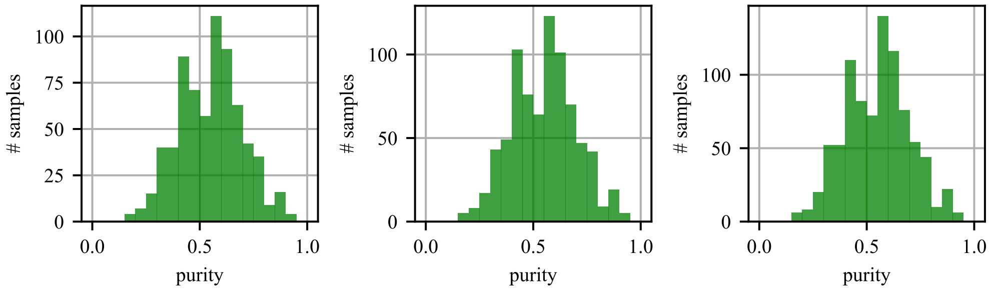


(a) training (b) validation (c) test

Figure 3: **KIRC cohort:** Genomic tumor purity histograms for training, validation, and test sets.

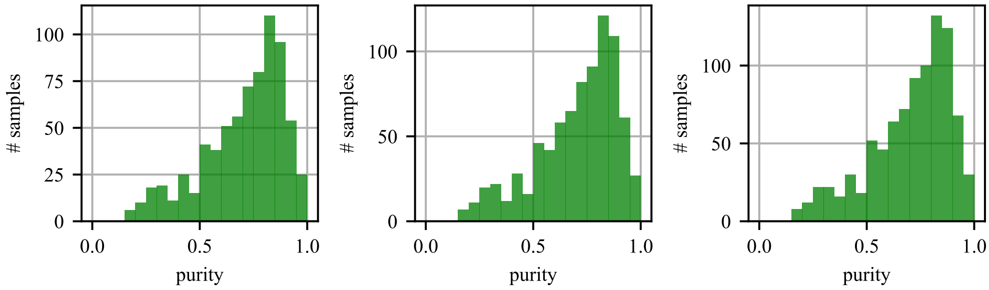


(a) training (b) validation (c) test

Figure 4: **LGG cohort:** Genomic tumor purity histograms for training, validation, and test sets.

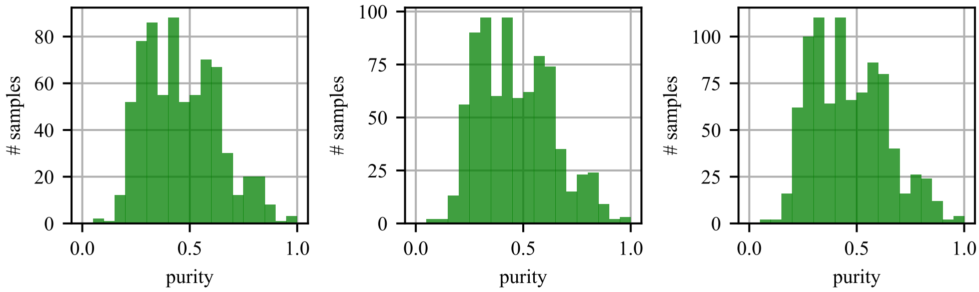


(a) training (b) validation (c) test

Figure 5: **LUAD cohort:** Genomic tumor purity histograms for training, validation, and test sets.

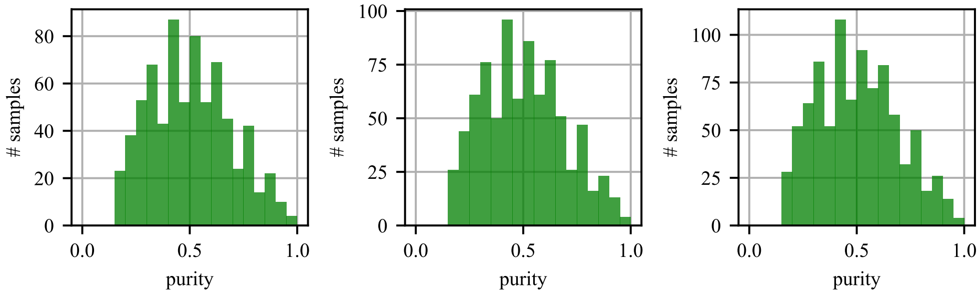


(a) training (b) validation (c) test

Figure 6: **LUSC cohort:** Genomic tumor purity histograms for training, validation, and test sets.

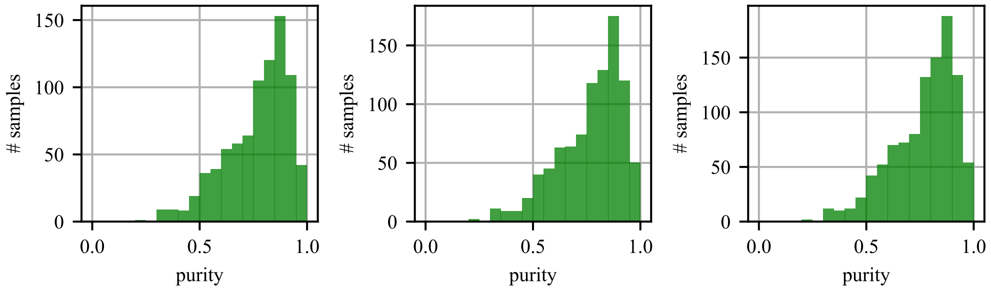


(a) training (b) validation (c) test

Figure 7: **OV cohort:** Genomic tumor purity histograms for training, validation, and test sets.

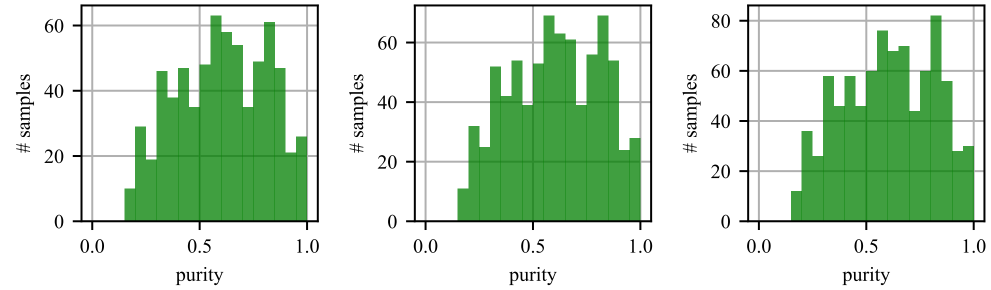


(a) training (b) validation (c) test

Figure 8: **PRAD cohort:** Genomic tumor purity histograms for training, validation, and test sets.

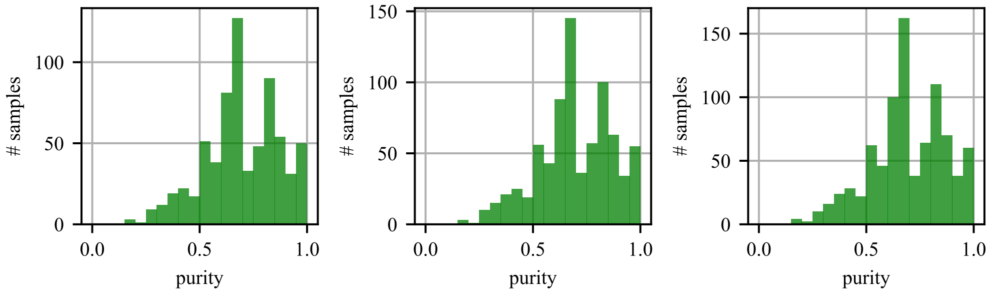


(a) training (b) validation (c) test

Figure 9: **THCA cohort:** Genomic tumor purity histograms for training, validation, and test sets.

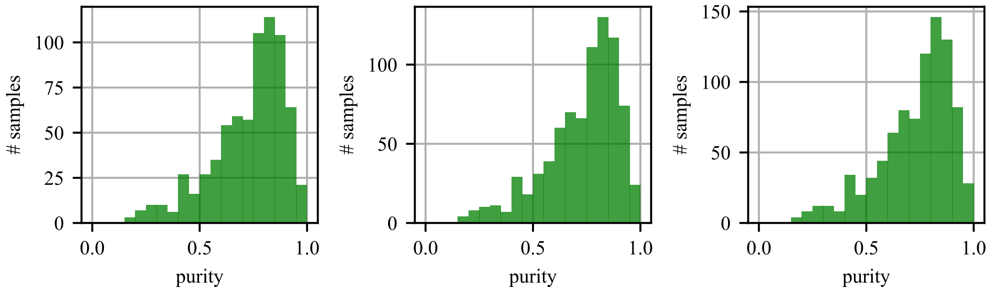


(a) training (b) validation (c) test

Figure 10: **UCEC cohort:** Genomic tumor purity histograms for training, validation, and test sets.

### 2 Tumor Purity Prediction Statistics in TCGA cohorts

#### 2.1 Scatter Plots and Correlation Analysis

Scatter plot of genomic tumor purity values obtained from ABSOLUTE and tumor purity predictions obtained from MIL model, and scatter plot of genomic tumor purity values obtained from ABSOLUTE and percent tumor nuclei values estimated by pathologists in each set are given for: BRCA cohort in Figure 11, GBM cohort in Figure 12, KIRC cohort in Figure 13, LGG cohort in Figure 14, LUAD cohort in Figure 15, LUSC cohort in Figure 16, OV cohort in Figure 17, PRAD cohort in Figure 18, THCA cohort in Figure 19, UCEC cohort in Figure 20.

Moreover, Spearman's correlation coefficients between genomic tumor purity obtained from ABSOLUTE and tumor purity prediction obtained from MIL model in the training, validation, and test sets for all samples and tumor samples only are presented for: BRCA cohort in Table 11, GBM cohort in Table 12, KIRC cohort in Table 13, LGG cohort in Table 14, LUAD cohort in Table 15, LUSC cohort in Table 16, OV cohort in Table 17, PRAD cohort in Table 18, THCA cohort in Table 19, UCEC cohort in Table 20.

We also conducted statistical tests on correlation coefficients to compare: (i) our MIL models' predictions and (ii) pathologists' percent tumor nuclei estimates. We used the Fisher's z transformation-based method of Meng et al. [2]. We compared two methods only when there was a significant correlation for both methods in a cohort. We summarized the results of statistical tests in Table 21.

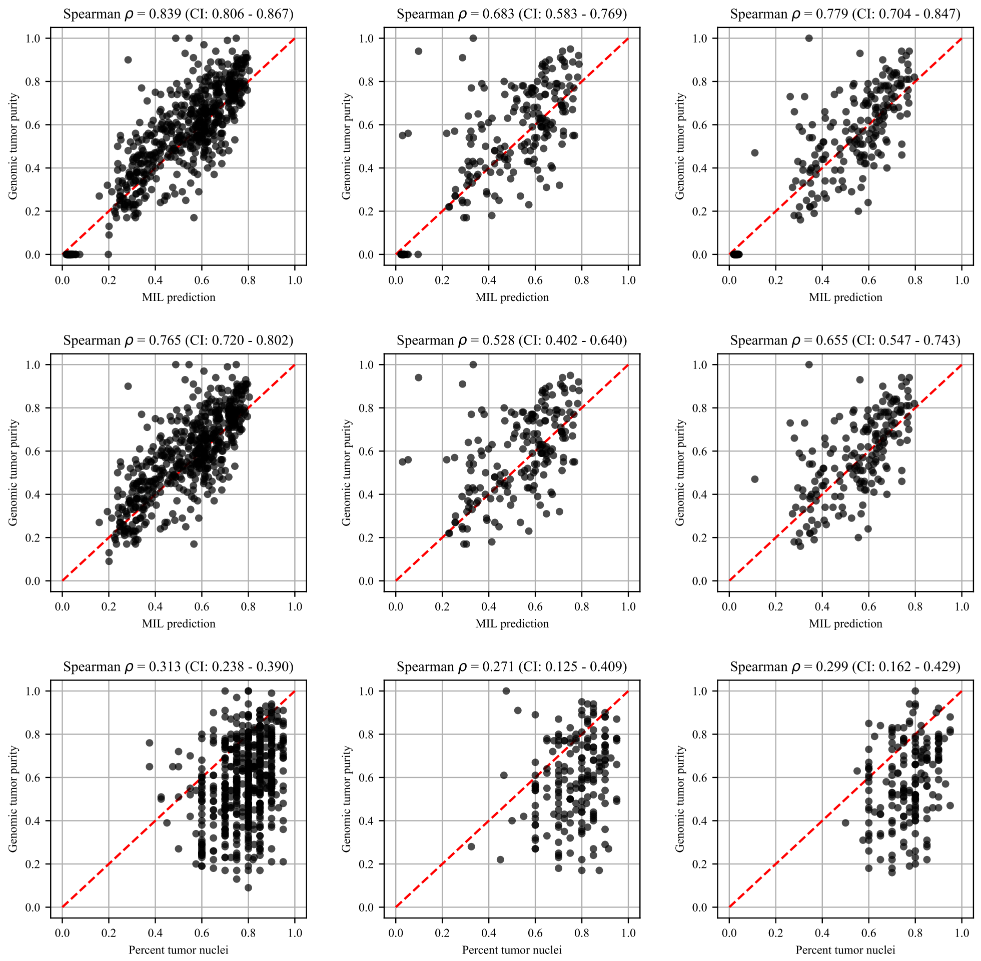


(a) Training (b) Validation (c) Test

Figure 11: **BRCA cohort.** Scatter plot of genomic tumor purity obtained from ABSOLUTE and tumor purity prediction obtained from MIL model for all samples (Top) and for tumor samples only (Middle). Scatter plot of genomic tumor purity obtained from ABSOLUTE and percent tumor nuclei estimated by pathologists (Bottom). Diagonal red dotted lines show the y=x line.

Table 11: **BRCA cohort.** Spearman's correlation coefficients between genomic tumor purity values and MIL predictions ($\rho_{mil}$) and genomic tumor purity values and pathologists' percent tumor nuclei estimates ($\rho_{path}$) are calculated in the training, validation, and test sets. Correlation coefficients together with calculated p-values ($P_{\rho_{mil}}$ and $P_{\rho_{path}}$) and 95% confidence intervals ($CI_{\rho_{mil}}$ and $CI_{\rho_{path}}$) are presented for all samples and tumor samples only.

| | MIL prediction | | | | | | Pathologist's estimate | | |
|---|---|---|---|---|---|---|---|---|---|
| | All samples | | | Tumor samples only | | | Tumor samples only | | |
| | $\rho_{mil}$ | $P_{\rho_{mil}}$ | $CI_{\rho_{mil}}$ | $\rho_{mil}$ | $P_{\rho_{mil}}$ | $CI_{\rho_{mil}}$ | $\rho_{path}$ | $P_{\rho_{path}}$ | $CI_{\rho_{path}}$ |
| train | 0.839 | 2.4e-169 | 0.806 - 0.867 | 0.765 | 1.2e-108 | 0.720 - 0.802 | 0.313 | 3.3e-14 | 0.238 - 0.390 |
| valid | 0.683 | 1.8e-30 | 0.583 - 0.769 | 0.528 | 1.1e-14 | 0.402 - 0.640 | 0.271 | 1.9e-04 | 0.125 - 0.409 |
| test | 0.779 | 5.2e-45 | 0.704 - 0.847 | 0.655 | 4.6e-24 | 0.547 - 0.743 | 0.299 | 3.6e-05 | 0.162 - 0.429 |

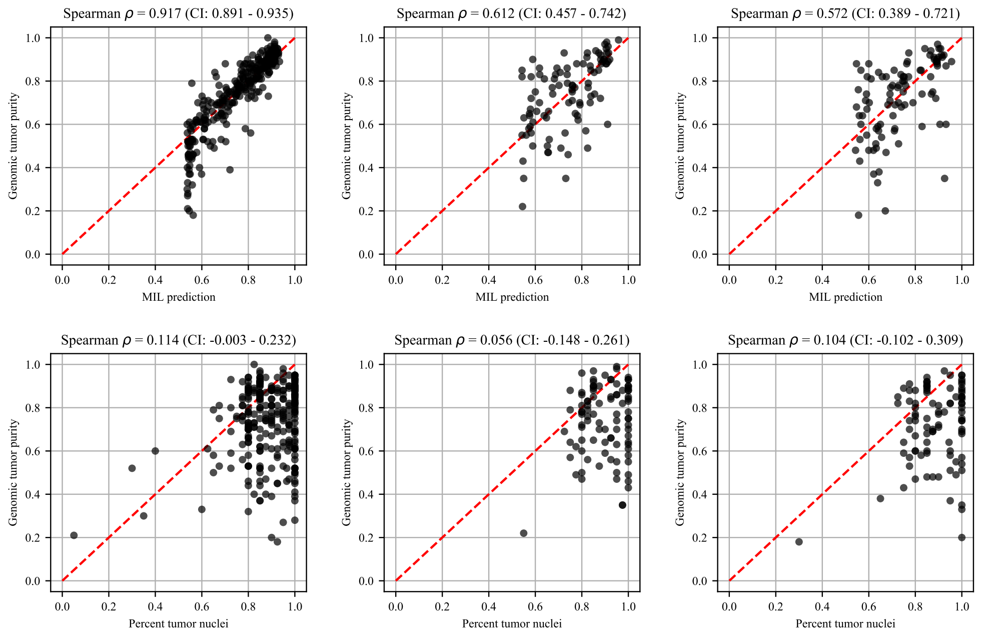


(a) Training (b) Validation (c) Test

Figure 12: **GBM cohort.** Scatter plot of genomic tumor purity obtained from ABSOLUTE and tumor purity prediction obtained from MIL model for tumor samples only (Top). Scatter plot of genomic tumor purity obtained from ABSOLUTE and percent tumor nuclei estimated by pathologists (Bottom). Diagonal red dotted lines show the y=x line. Note that there are no normal slides in the GBM cohort.

Table 12: **GBM cohort.** Spearman's correlation coefficients between genomic tumor purity values and MIL predictions ($\rho_{mil}$) and genomic tumor purity values and pathologists' percent tumor nuclei estimates ($\rho_{path}$) are calculated in the training, validation, and test sets. Correlation coefficients together with calculated p-values ($P_{\rho_{mil}}$ and $P_{\rho_{path}}$) and 95% confidence intervals ($CI_{\rho_{mil}}$ and $CI_{\rho_{path}}$) are presented for tumor samples only. Note that there are no normal slides in the GBM cohort.

| | MIL prediction | | | Pathologist's estimate | | |
|---|---|---|---|---|---|---|
| | Tumor samples only | | | Tumor samples only | | |
| | $\rho_{mil}$ | $P_{\rho_{mil}}$ | $CI_{\rho_{mil}}$ | $\rho_{path}$ | $P_{\rho_{path}}$ | $CI_{\rho_{path}}$ |
| train | 0.917 | 9.7e-115 | 0.891 - 0.935 | 0.114 | 5.4e-02 | -0.003 - 0.232 |
| valid | 0.612 | 4.5e-11 | 0.457 - 0.742 | 0.056 | 5.9e-01 | -0.148 - 0.261 |
| test | 0.572 | 1.7e-09 | 0.389 - 0.721 | 0.104 | 3.2e-01 | -0.102 - 0.309 |

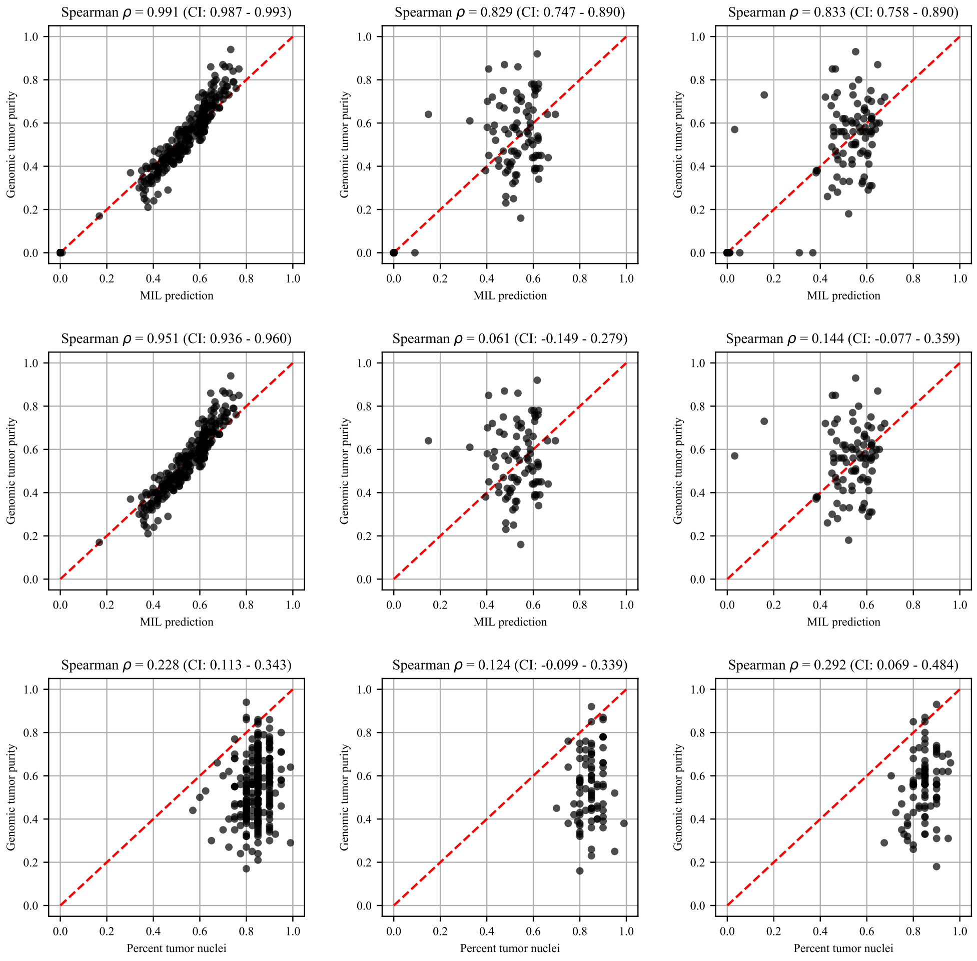


(a) Training (b) Validation (c) Test

Figure 13: **KIRC cohort.** Scatter plot of genomic tumor purity obtained from ABSOLUTE and tumor purity prediction obtained from MIL model for all samples (Top) and for tumor samples only (Middle). Scatter plot of genomic tumor purity obtained from ABSOLUTE and percent tumor nuclei estimated by pathologists (Bottom). Diagonal red dotted lines show the y=x line.

Table 13: **KIRC cohort.** Spearman's correlation coefficients between genomic tumor purity values and MIL predictions ($\rho_{mil}$) and genomic tumor purity values and pathologists' percent tumor nuclei estimates ($\rho_{path}$) are calculated in the training, validation, and test sets. Correlation coefficients together with calculated p-values ($P_{\rho_{mil}}$ and $P_{\rho_{path}}$) and 95% confidence intervals ($CI_{\rho_{mil}}$ and $CI_{\rho_{path}}$) are presented for all samples and tumor samples only.

| | MIL prediction | | | | | | Pathologist's estimate | | |
|---|---|---|---|---|---|---|---|---|---|
| | All samples | | | Tumor samples only | | | Tumor samples only | | |
| | $\rho_{mil}$ | $P_{\rho_{mil}}$ | $CI_{\rho_{mil}}$ | $\rho_{mil}$ | $P_{\rho_{mil}}$ | $CI_{\rho_{mil}}$ | $\rho_{path}$ | $P_{\rho_{path}}$ | $CI_{\rho_{path}}$ |
| train | 0.991 | 0.0e+00 | 0.987 - 0.993 | 0.951 | 4.3e-134 | 0.936 - 0.960 | 0.228 | 2.0e-04 | 0.113 - 0.343 |
| valid | 0.829 | 1.2e-40 | 0.747 - 0.890 | 0.061 | 5.8e-01 | -0.149 - 0.279 | 0.124 | 2.6e-01 | -0.099 - 0.339 |
| test | 0.833 | 5.5e-43 | 0.758 - 0.890 | 0.144 | 1.8e-01 | -0.077 - 0.359 | 0.292 | 5.5e-03 | 0.069 - 0.484 |

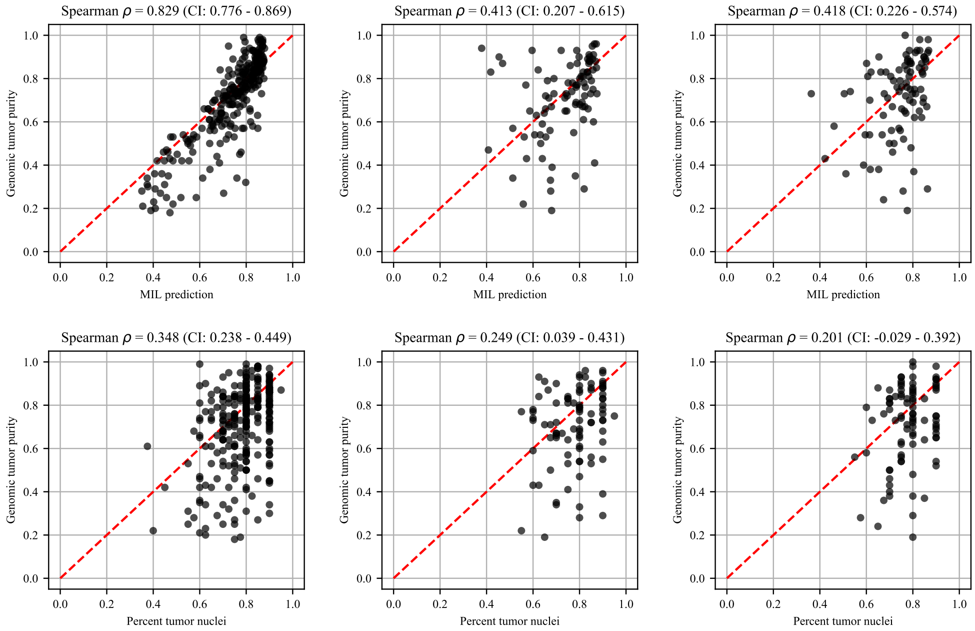


(a) Training (b) Validation (c) Test

Figure 14: **LGG cohort.** Scatter plot of genomic tumor purity obtained from ABSOLUTE and tumor purity prediction obtained from MIL model for tumor samples only (Top). Scatter plot of genomic tumor purity obtained from ABSOLUTE and percent tumor nuclei estimated by pathologists (Bottom). Diagonal red dotted lines show the y=x line. Note that there are no normal slides in the LGG cohort.

Table 14: **LGG cohort.** Spearman's correlation coefficients between genomic tumor purity values and MIL predictions ($\rho_{mil}$) and genomic tumor purity values and pathologists' percent tumor nuclei estimates ($\rho_{path}$) are calculated in the training, validation, and test sets. Correlation coefficients together with calculated p-values ($P_{\rho_{mil}}$ and $P_{\rho_{path}}$) and 95% confidence intervals ($CI_{\rho_{mil}}$ and $CI_{\rho_{path}}$) are presented for tumor samples only. Note that there are no normal slides in the LGG cohort.

| | MIL prediction | | | Pathologist's estimate | | |
|---|---|---|---|---|---|---|
| | Tumor samples only | | | Tumor samples only | | |
| | $\rho_{mil}$ | $P_{\rho_{mil}}$ | $CI_{\rho_{mil}}$ | $\rho_{path}$ | $P_{\rho_{path}}$ | $CI_{\rho_{path}}$ |
| train | 0.829 | 2.7e-70 | 0.776 - 0.869 | 0.348 | 3.5e-09 | 0.238 - 0.449 |
| valid | 0.413 | 4.7e-05 | 0.207 - 0.615 | 0.249 | 1.7e-02 | 0.039 - 0.431 |
| test | 0.418 | 4.1e-05 | 0.226 - 0.574 | 0.201 | 5.7e-02 | -0.029 - 0.392 |

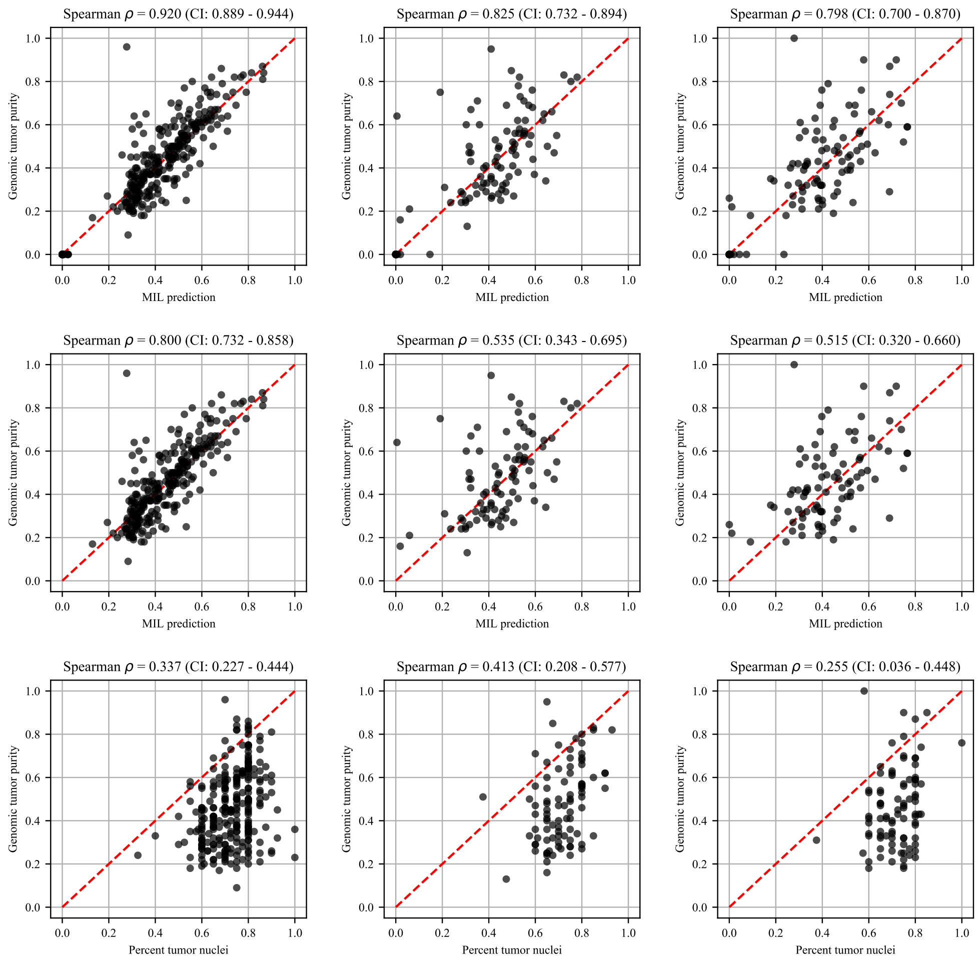


(a) Training (b) Validation (c) Test

Figure 15: **LUAD cohort.** Scatter plot of genomic tumor purity obtained from ABSOLUTE and tumor purity prediction obtained from MIL model for all samples (Top) and for tumor samples only (Middle). Scatter plot of genomic tumor purity obtained from ABSOLUTE and percent tumor nuclei estimated by pathologists (Bottom). Diagonal red dotted lines show the y=x line.

Table 15: **LUAD cohort.** Spearman's correlation coefficients between genomic tumor purity values and MIL predictions ($\rho_{mil}$) and genomic tumor purity values and pathologists' percent tumor nuclei estimates ($\rho_{path}$) are calculated in the training, validation, and test sets. Correlation coefficients together with calculated p-values ($P_{\rho_{mil}}$ and $P_{\rho_{path}}$) and 95% confidence intervals ($CI_{\rho_{mil}}$ and $CI_{\rho_{path}}$) are presented for all samples and tumor samples only.

| | MIL prediction | | | | | | Pathologist's estimate | | |
|---|---|---|---|---|---|---|---|---|---|
| | All samples | | | Tumor samples only | | | Tumor samples only | | |
| | $\rho_{mil}$ | $P_{\rho_{mil}}$ | $CI_{\rho_{mil}}$ | $\rho_{mil}$ | $P_{\rho_{mil}}$ | $CI_{\rho_{mil}}$ | $\rho_{path}$ | $P_{\rho_{path}}$ | $CI_{\rho_{path}}$ |
| train | 0.920 | 2.4e-150 | 0.889 - 0.944 | 0.800 | 1.3e-60 | 0.732 - 0.858 | 0.337 | 1.7e-08 | 0.227 - 0.444 |
| valid | 0.825 | 8.5e-33 | 0.732 - 0.894 | 0.535 | 5.7e-08 | 0.343 - 0.695 | 0.413 | 5.2e-05 | 0.208 - 0.577 |
| test | 0.798 | 2.2e-28 | 0.700 - 0.870 | 0.515 | 2.1e-07 | 0.320 - 0.660 | 0.255 | 1.5e-02 | 0.036 - 0.448 |

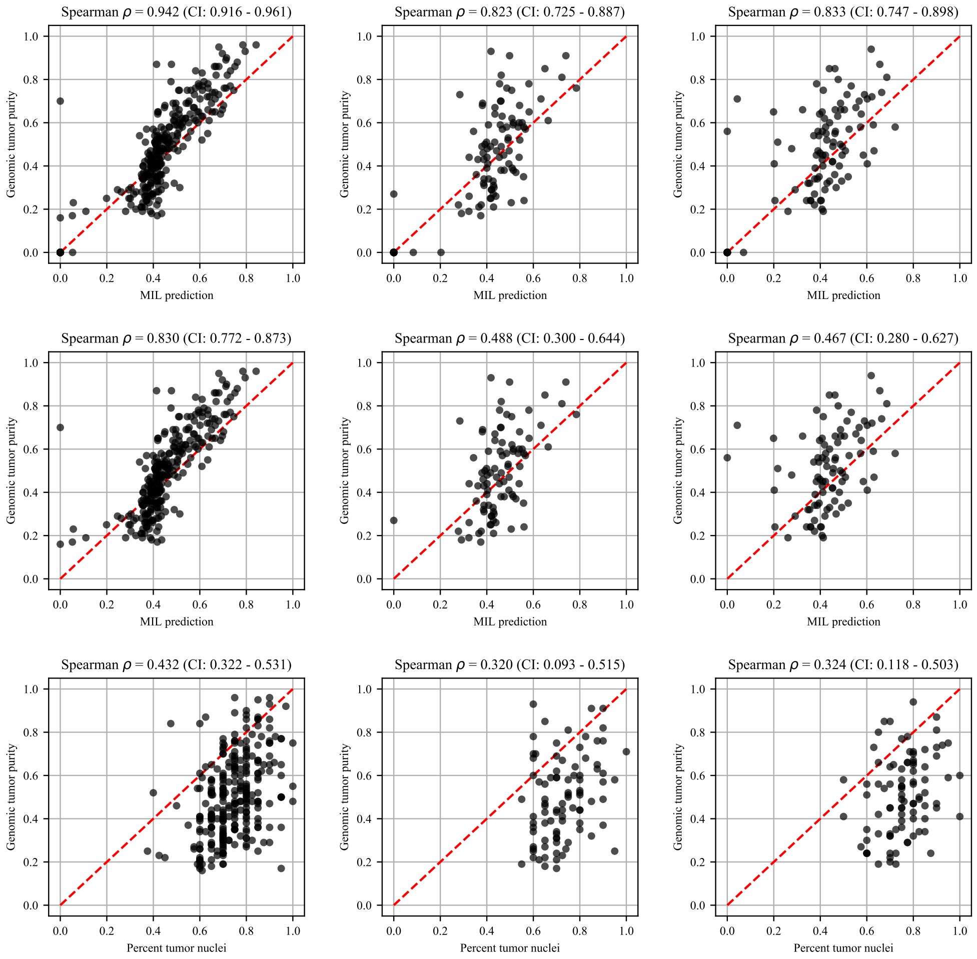


(a) Training (b) Validation (c) Test

Figure 16: **LUSC cohort.** Scatter plot of genomic tumor purity obtained from ABSOLUTE and tumor purity prediction obtained from MIL model for all samples (Top) and for tumor samples only (Middle). Scatter plot of genomic tumor purity obtained from ABSOLUTE and percent tumor nuclei estimated by pathologists (Bottom). Diagonal red dotted lines show the y=x line.

Table 16: **LUSC cohort.** Spearman's correlation coefficients between genomic tumor purity values and MIL predictions ($\rho_{mil}$) and genomic tumor purity values and pathologists' percent tumor nuclei estimates ($\rho_{path}$) are calculated in the training, validation, and test sets. Correlation coefficients together with calculated p-values ($P_{\rho_{mil}}$ and $P_{\rho_{path}}$) and 95% confidence intervals ($CI_{\rho_{mil}}$ and $CI_{\rho_{path}}$) are presented for all samples and tumor samples only.

| | MIL prediction | | | | | | Pathologist's estimate | | |
|---|---|---|---|---|---|---|---|---|---|
| | All samples | | | Tumor samples only | | | Tumor samples only | | |
| | $\rho_{mil}$ | $P_{\rho_{mil}}$ | $CI_{\rho_{mil}}$ | $\rho_{mil}$ | $P_{\rho_{mil}}$ | $CI_{\rho_{mil}}$ | $\rho_{path}$ | $P_{\rho_{path}}$ | $CI_{\rho_{path}}$ |
| train | 0.942 | 1.7e-192 | 0.916 - 0.961 | 0.830 | 1.2e-70 | 0.772 - 0.873 | 0.432 | 7.4e-14 | 0.322 - 0.531 |
| valid | 0.823 | 2.0e-33 | 0.725 - 0.887 | 0.488 | 1.1e-06 | 0.300 - 0.644 | 0.320 | 2.1e-03 | 0.093 - 0.515 |
| test | 0.833 | 1.6e-36 | 0.747 - 0.898 | 0.467 | 3.5e-06 | 0.280 - 0.627 | 0.324 | 1.8e-03 | 0.118 - 0.503 |

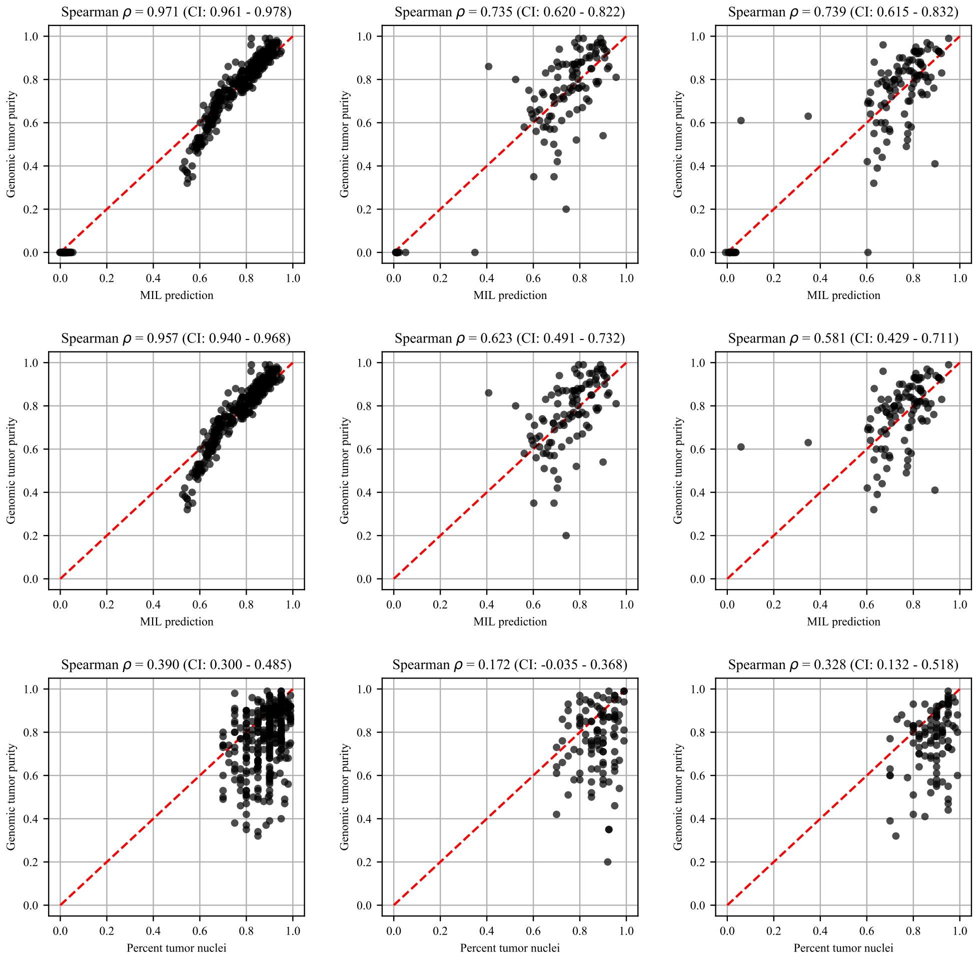


(a) Training (b) Validation (c) Test

Figure 17: **OV cohort.** Scatter plot of genomic tumor purity obtained from ABSOLUTE and tumor purity prediction obtained from MIL model for all samples (Top) and for tumor samples only (Middle). Scatter plot of genomic tumor purity obtained from ABSOLUTE and percent tumor nuclei estimated by pathologists (Bottom). Diagonal red dotted lines show the y=x line.

Table 17: **OV cohort.** Spearman's correlation coefficients between genomic tumor purity values and MIL predictions ($\rho_{mil}$) and genomic tumor purity values and pathologists' percent tumor nuclei estimates ($\rho_{path}$) are calculated in the training, validation, and test sets. Correlation coefficients together with calculated p-values ($P_{\rho_{mil}}$ and $P_{\rho_{path}}$) and 95% confidence intervals ($CI_{\rho_{mil}}$ and $CI_{\rho_{path}}$) are presented for all samples and tumor samples only.

| | MIL prediction | | | | | | Pathologist's estimate | | |
|---|---|---|---|---|---|---|---|---|---|
| | All samples | | | Tumor samples only | | | Tumor samples only | | |
| | $\rho_{mil}$ | $P_{\rho_{mil}}$ | $CI_{\rho_{mil}}$ | $\rho_{mil}$ | $P_{\rho_{mil}}$ | $CI_{\rho_{mil}}$ | $\rho_{path}$ | $P_{\rho_{path}}$ | $CI_{\rho_{path}}$ |
| train | 0.971 | 1.7e-227 | 0.961 - 0.978 | 0.957 | 4.9e-167 | 0.940 - 0.968 | 0.390 | 1.0e-12 | 0.300 - 0.485 |
| valid | 0.735 | 5.4e-21 | 0.620 - 0.822 | 0.623 | 2.0e-12 | 0.491 - 0.732 | 0.172 | 8.3e-02 | -0.035 - 0.368 |
| test | 0.739 | 4.0e-22 | 0.615 - 0.832 | 0.581 | 1.3e-10 | 0.429 - 0.711 | 0.328 | 7.1e-04 | 0.132 - 0.518 |

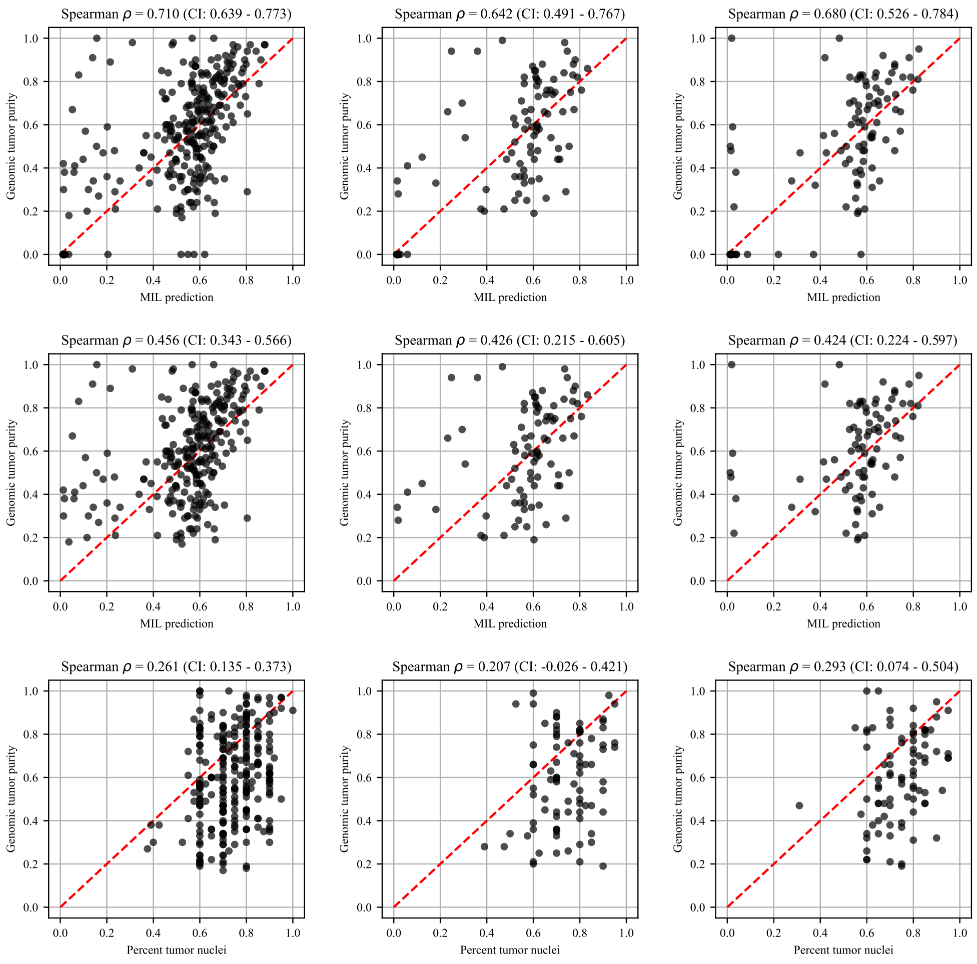


(a) Training (b) Validation (c) Test

Figure 18: **PRAD cohort.** Scatter plot of genomic tumor purity obtained from ABSOLUTE and tumor purity prediction obtained from MIL model for all samples (Top) and for tumor samples only (Middle). Scatter plot of genomic tumor purity obtained from ABSOLUTE and percent tumor nuclei estimated by pathologists (Bottom). Diagonal red dotted lines show the y=x line.

Table 18: **PRAD cohort.** Spearman's correlation coefficients between genomic tumor purity values and MIL predictions ($\rho_{mil}$) and genomic tumor purity values and pathologists' percent tumor nuclei estimates ($\rho_{path}$) are calculated in the training, validation, and test sets. Correlation coefficients together with calculated p-values ($P_{\rho_{mil}}$ and $P_{\rho_{path}}$) and 95% confidence intervals ($CI_{\rho_{mil}}$ and $CI_{\rho_{path}}$) are presented for all samples and tumor samples only.

| | MIL prediction | | | | | | Pathologist's estimate | | |
|---|---|---|---|---|---|---|---|---|---|
| | All samples | | | Tumor samples only | | | Tumor samples only | | |
| | $\rho_{mil}$ | $P_{\rho_{mil}}$ | $CI_{\rho_{mil}}$ | $\rho_{mil}$ | $P_{\rho_{mil}}$ | $CI_{\rho_{mil}}$ | $\rho_{path}$ | $P_{\rho_{path}}$ | $CI_{\rho_{path}}$ |
| train | 0.710 | 8.5e-52 | 0.639 - 0.773 | 0.456 | 1.1e-14 | 0.343 - 0.566 | 0.261 | 2.2e-05 | 0.135 - 0.373 |
| valid | 0.642 | 6.0e-13 | 0.491 - 0.767 | 0.426 | 4.8e-05 | 0.215 - 0.605 | 0.207 | 5.8e-02 | -0.026 - 0.421 |
| test | 0.680 | 4.1e-16 | 0.526 - 0.784 | 0.424 | 5.3e-05 | 0.224 - 0.597 | 0.293 | 6.5e-03 | 0.074 - 0.504 |

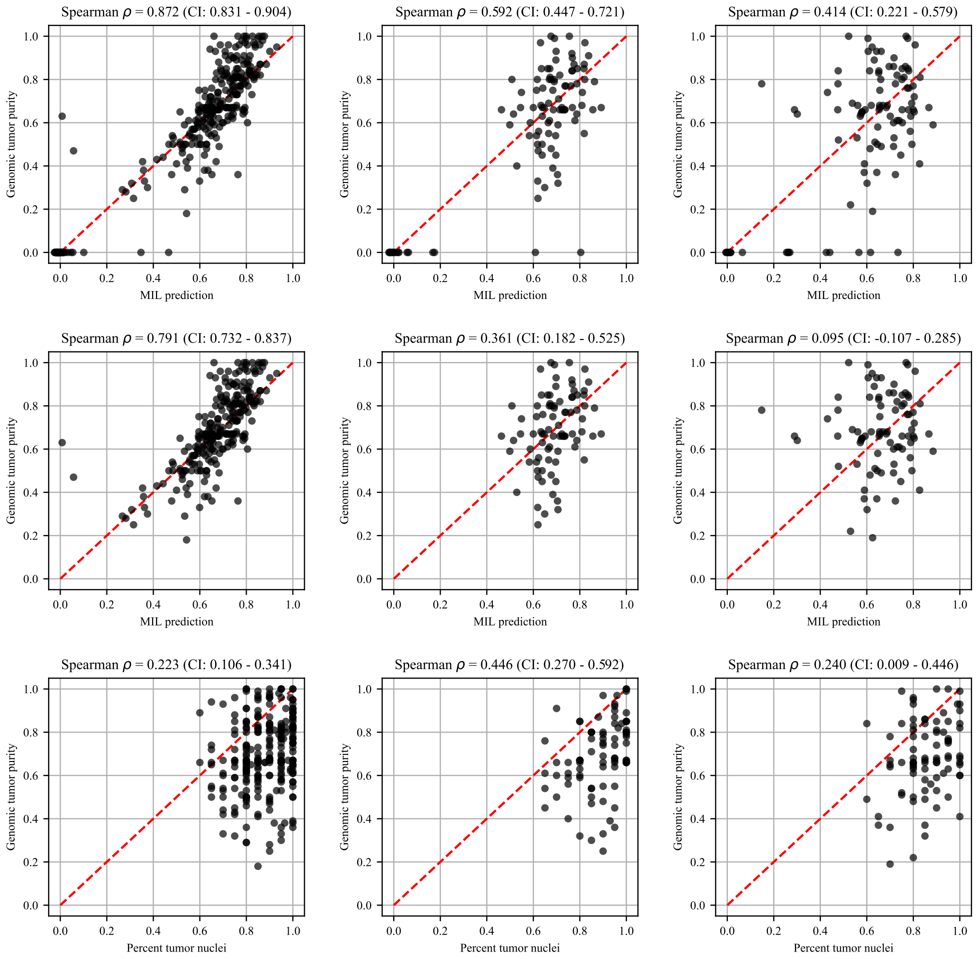


(a) Training (b) Validation (c) Test

Figure 19: **THCA cohort.** Scatter plot of genomic tumor purity obtained from ABSOLUTE and tumor purity prediction obtained from MIL model for all samples (Top) and for tumor samples only (Middle). Scatter plot of genomic tumor purity obtained from ABSOLUTE and percent tumor nuclei estimated by pathologists (Bottom). Diagonal red dotted lines show the y=x line.

Table 19: **THCA cohort.** Spearman's correlation coefficients between genomic tumor purity values and MIL predictions ($\rho_{mil}$) and genomic tumor purity values and pathologists' percent tumor nuclei estimates ($\rho_{path}$) are calculated in the training, validation, and test sets. Correlation coefficients together with calculated p-values ($P_{\rho_{mil}}$ and $P_{\rho_{path}}$) and 95% confidence intervals ($CI_{\rho_{mil}}$ and $CI_{\rho_{path}}$) are presented for all samples and tumor samples only.

| | MIL prediction | | | | | | Pathologist's estimate | | |
|---|---|---|---|---|---|---|---|---|---|
| | All samples | | | Tumor samples only | | | Tumor samples only | | |
| | $\rho_{mil}$ | $P_{\rho_{mil}}$ | $CI_{\rho_{mil}}$ | $\rho_{mil}$ | $P_{\rho_{mil}}$ | $CI_{\rho_{mil}}$ | $\rho_{path}$ | $P_{\rho_{path}}$ | $CI_{\rho_{path}}$ |
| train | 0.872 | 2.3e-96 | 0.831 - 0.904 | 0.791 | 1.2e-56 | 0.732 - 0.837 | 0.223 | 3.1e-04 | 0.106 - 0.341 |
| valid | 0.592 | 4.4e-11 | 0.447 - 0.721 | 0.361 | 7.0e-04 | 0.182 - 0.525 | 0.446 | 1.9e-05 | 0.270 - 0.592 |
| test | 0.414 | 1.6e-05 | 0.221 - 0.579 | 0.095 | 3.9e-01 | -0.107 - 0.285 | 0.240 | 2.7e-02 | 0.009 - 0.446 |

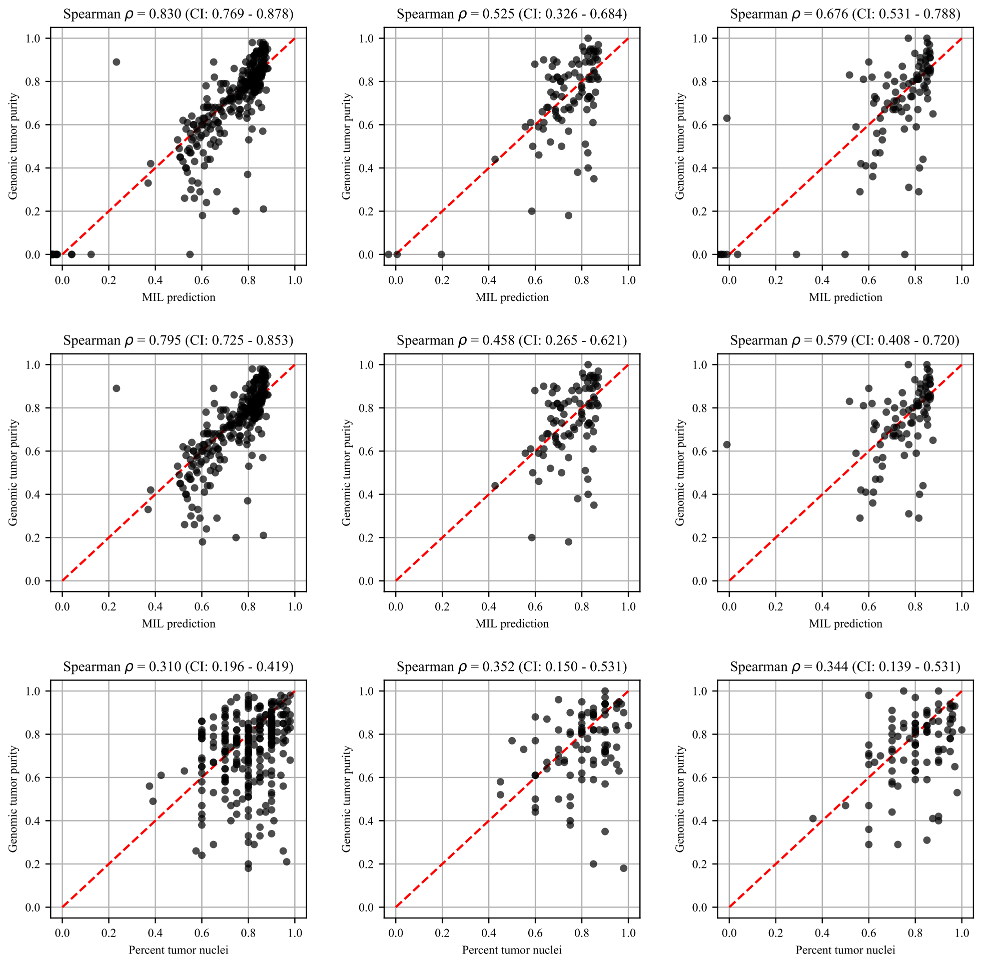


(a) Training (b) Validation (c) Test

Figure 20: **UCEC cohort.** Scatter plot of genomic tumor purity obtained from ABSOLUTE and tumor purity prediction obtained from MIL model for all samples (Top) and for tumor samples only (Middle). Scatter plot of genomic tumor purity obtained from ABSOLUTE and percent tumor nuclei estimated by pathologists (Bottom). Diagonal red dotted lines show the y=x line.

Table 20: **UCEC cohort.** Spearman's correlation coefficients between genomic tumor purity values and MIL predictions ($\rho_{mil}$) and genomic tumor purity values and pathologists' percent tumor nuclei estimates ($\rho_{path}$) are calculated in the training, validation, and test sets. Correlation coefficients together with calculated p-values ($P_{\rho_{mil}}$ and $P_{\rho_{path}}$) and 95% confidence intervals ($CI_{\rho_{mil}}$ and $CI_{\rho_{path}}$) are presented for all samples and tumor samples only.

| | MIL prediction | | | | | | Pathologist's estimate | | |
|---|---|---|---|---|---|---|---|---|---|
| | All samples | | | Tumor samples only | | | Tumor samples only | | |
| | $\rho_{mil}$ | $P_{\rho_{mil}}$ | $CI_{\rho_{mil}}$ | $\rho_{mil}$ | $P_{\rho_{mil}}$ | $CI_{\rho_{mil}}$ | $\rho_{path}$ | $P_{\rho_{path}}$ | $CI_{\rho_{path}}$ |
| train | 0.830 | 1.3e-74 | 0.769 - 0.878 | 0.795 | 4.4e-60 | 0.725 - 0.853 | 0.310 | 2.1e-07 | 0.196 - 0.419 |
| valid | 0.525 | 5.7e-08 | 0.326 - 0.684 | 0.458 | 5.5e-06 | 0.265 - 0.621 | 0.352 | 6.8e-04 | 0.150 - 0.531 |
| test | 0.676 | 1.6e-14 | 0.531 - 0.788 | 0.579 | 2.7e-09 | 0.408 - 0.720 | 0.344 | 9.8e-04 | 0.139 - 0.531 |

Table 21: **Comparison of methods based on Spearman's correlation coefficients in the test sets of different cohorts.** Spearman's correlation coefficients between genomic tumor purity values and MIL predictions ($\rho_{mil}$) and genomic tumor purity values and pathologists' percent tumor nuclei estimates ($\rho_{path}$) in the test sets of different cohorts are calculated for only the tumor samples. Then, they are compared using the method in Meng et al. [2]. Spearman's correlation coefficients together with calculated p-values ($P_{\rho_{mil}}$ and $P_{\rho_{path}}$) and 95% confidence intervals ($CI_{\rho_{mil}}$ and $CI_{\rho_{path}}$) and calculated p-values in statistical tests ($P_{comp}$) are presented. Note that if the calculated correlation in any method is not significant (i.e., $P_{\rho_{mil}} > 5.0e-02$ or $P_{\rho_{path}} > 5.0e-02$), the statistical test is not conducted. It is indicated by 'x'. The best methods are highlighted in bold.

| | MIL prediction | | | Pathologist's estimate | | | Comparison |
|---|---|---|---|---|---|---|---|
| | $\rho_{mil}$ | $P_{\rho_{mil}}$ | $CI_{\rho_{mil}}$ | $\rho_{path}$ | $P_{\rho_{path}}$ | $CI_{\rho_{path}}$ | $P_{comp}$ |
| BRCA | **0.655** | **4.6e-24** | **0.547 - 0.743** | 0.299 | 3.6e-05 | 0.162 - 0.429 | **1.4e-07** |
| GBM | 0.572 | 1.7e-09 | 0.389 - 0.721 | 0.104 | 3.2e-01 | -0.102 - 0.309 | x |
| KIRC | 0.144 | 1.8e-01 | -0.077 - 0.359 | 0.292 | 5.5e-03 | 0.069 - 0.484 | x |
| LGG | 0.418 | 4.1e-05 | 0.226 - 0.574 | 0.201 | 5.7e-02 | -0.029 - 0.392 | x |
| LUAD | **0.515** | **2.1e-07** | **0.320 - 0.660** | 0.255 | 1.5e-02 | 0.036 - 0.448 | **1.2e-02** |
| LUSC | 0.467 | 3.5e-06 | 0.280 - 0.627 | 0.324 | 1.8e-03 | 0.118 - 0.503 | 1.7e-01 |
| OV | **0.581** | **1.3e-10** | **0.429 - 0.711** | 0.328 | 7.1e-04 | 0.132 - 0.518 | **9.4e-03** |
| PRAD | 0.424 | 5.3e-05 | 0.224 - 0.597 | 0.293 | 6.5e-03 | 0.074 - 0.504 | 2.0e-01 |
| THCA | 0.095 | 3.9e-01 | -0.107 - 0.285 | 0.240 | 2.7e-02 | 0.009 - 0.446 | x |
| UCEC | **0.579** | **2.7e-09** | **0.408 - 0.720** | 0.344 | 9.8e-04 | 0.139 - 0.531 | **2.6e-02** |

#### 2.2 Absolute Error Analysis

The results of absolute error analyses between genomic tumor purity values and MIL predictions in the training, validation, and test sets for tumor samples only are presented for: BRCA cohort in Table 22, GBM cohort in Table 23, KIRC cohort in Table 24, LGG cohort in Table 25, LUAD cohort in Table 26, LUSC cohort in Table 27, OV cohort in Table 28, PRAD cohort in Table 29, THCA cohort in Table 30, UCEC cohort in Table 31.

Similar to our comparison in correlation analyses, we compared two methods based on absolute errors in the test sets of different cohorts. We used the Wilcoxon signed-rank test [3] on absolute error values for tumor samples in the test sets. We summarized the results of statistical tests in Table 32.

Table 22: **BRCA cohort.** Absolute errors between genomic tumor purity values and MIL predictions ($e_{mil}$) and genomic tumor purity values and pathologists' percent tumor nuclei estimates ($e_{path}$) in the training, validation, and test sets are calculated using only the tumor samples. Mean absolute errors ($\mu_{e_{mil}}$ and $\mu_{e_{path}}$) together with standard deviations ($\sigma_{e_{mil}}$ and $\sigma_{e_{path}}$) and median absolute errors ($m_{e_{mil}}$ and $m_{e_{path}}$) together with interquartile ranges ($IQR_{e_{mil}}$ and $IQR_{e_{path}}$) are presented.

| | MIL prediction | | | | Pathologist's estimate | | | |
|---|---|---|---|---|---|---|---|---|
| | $\mu_{e_{mil}}$ | $\sigma_{e_{mil}}$ | $m_{e_{mil}}$ | $IQR_{e_{mil}}$ | $\mu_{e_{path}}$ | $\sigma_{e_{path}}$ | $m_{e_{path}}$ | $IQR_{e_{path}}$ |
| training | 0.097 | 0.083 | 0.076 | 0.035 - 0.134 | 0.219 | 0.155 | 0.190 | 0.090 - 0.320 |
| validation | 0.139 | 0.128 | 0.104 | 0.043 - 0.200 | 0.215 | 0.157 | 0.190 | 0.090 - 0.315 |
| test | 0.116 | 0.097 | 0.104 | 0.043 - 0.159 | 0.220 | 0.147 | 0.200 | 0.105 - 0.310 |

Table 23: **GBM cohort.** Absolute errors between genomic tumor purity values and MIL predictions ($e_{mil}$) and genomic tumor purity values and pathologists' percent tumor nuclei estimates ($e_{path}$) in the training, validation, and test sets are calculated using only the tumor samples. Mean absolute errors ($\mu_{e_{mil}}$ and $\mu_{e_{path}}$) together with standard deviations ($\sigma_{e_{mil}}$ and $\sigma_{e_{path}}$) and median absolute errors ($m_{e_{mil}}$ and $m_{e_{path}}$) together with interquartile ranges ($IQR_{e_{mil}}$ and $IQR_{e_{path}}$) are presented.

| | MIL prediction | | | | Pathologist's estimate | | | |
|---|---|---|---|---|---|---|---|---|
| | $\mu_{e_{mil}}$ | $\sigma_{e_{mil}}$ | $m_{e_{mil}}$ | $IQR_{e_{mil}}$ | $\mu_{e_{path}}$ | $\sigma_{e_{path}}$ | $m_{e_{path}}$ | $IQR_{e_{path}}$ |
| training | 0.055 | 0.063 | 0.032 | 0.018 - 0.065 | 0.179 | 0.157 | 0.130 | 0.070 - 0.250 |
| validation | 0.101 | 0.092 | 0.079 | 0.023 - 0.143 | 0.181 | 0.154 | 0.130 | 0.060 - 0.265 |
| test | 0.113 | 0.106 | 0.074 | 0.046 - 0.142 | 0.195 | 0.158 | 0.145 | 0.080 - 0.260 |

Table 24: **KIRC cohort.** Absolute errors between genomic tumor purity values and MIL predictions ($e_{mil}$) and genomic tumor purity values and pathologists' percent tumor nuclei estimates ($e_{path}$) in the training, validation, and test sets are calculated using only the tumor samples. Mean absolute errors ($\mu_{e_{mil}}$ and $\mu_{e_{path}}$) together with standard deviations ($\sigma_{e_{mil}}$ and $\sigma_{e_{path}}$) and median absolute errors ($m_{e_{mil}}$ and $m_{e_{path}}$) together with interquartile ranges ($IQR_{e_{mil}}$ and $IQR_{e_{path}}$) are presented.

| | MIL prediction | | | | Pathologist's estimate | | | |
|---|---|---|---|---|---|---|---|---|
| | $\mu_{e_{mil}}$ | $\sigma_{e_{mil}}$ | $m_{e_{mil}}$ | $IQR_{e_{mil}}$ | $\mu_{e_{path}}$ | $\sigma_{e_{path}}$ | $m_{e_{path}}$ | $IQR_{e_{path}}$ |
| training | 0.045 | 0.038 | 0.038 | 0.018 - 0.060 | 0.293 | 0.138 | 0.290 | 0.185 - 0.385 |
| validation | 0.146 | 0.103 | 0.133 | 0.069 - 0.196 | 0.301 | 0.159 | 0.305 | 0.175 - 0.415 |
| test | 0.128 | 0.119 | 0.082 | 0.042 - 0.189 | 0.298 | 0.141 | 0.290 | 0.220 - 0.395 |

Table 25: **LGG cohort.** Absolute errors between genomic tumor purity values and MIL predictions ($e_{mil}$) and genomic tumor purity values and pathologists' percent tumor nuclei estimates ($e_{path}$) in the training, validation, and test sets are calculated using only the tumor samples. Mean absolute errors ($\mu_{e_{mil}}$ and $\mu_{e_{path}}$) together with standard deviations ($\sigma_{e_{mil}}$ and $\sigma_{e_{path}}$) and median absolute errors ($m_{e_{mil}}$ and $m_{e_{path}}$) together with interquartile ranges ($IQR_{e_{mil}}$ and $IQR_{e_{path}}$) are presented.

| | MIL prediction | | | | Pathologist's estimate | | | |
|---|---|---|---|---|---|---|---|---|
| | $\mu_{e_{mil}}$ | $\sigma_{e_{mil}}$ | $m_{e_{mil}}$ | $IQR_{e_{mil}}$ | $\mu_{e_{path}}$ | $\sigma_{e_{path}}$ | $m_{e_{path}}$ | $IQR_{e_{path}}$ |
| training | 0.075 | 0.079 | 0.049 | 0.023 - 0.101 | 0.142 | 0.130 | 0.100 | 0.040 - 0.210 |
| validation | 0.128 | 0.135 | 0.088 | 0.030 - 0.153 | 0.151 | 0.130 | 0.120 | 0.050 - 0.205 |
| test | 0.136 | 0.119 | 0.105 | 0.052 - 0.188 | 0.152 | 0.122 | 0.130 | 0.060 - 0.200 |

Table 26: **LUAD cohort.** Absolute errors between genomic tumor purity values and MIL predictions ($e_{mil}$) and genomic tumor purity values and pathologists' percent tumor nuclei estimates ($e_{path}$) in the training, validation, and test sets are calculated using only the tumor samples. Mean absolute errors ($\mu_{e_{mil}}$ and $\mu_{e_{path}}$) together with standard deviations ($\sigma_{e_{mil}}$ and $\sigma_{e_{path}}$) and median absolute errors ($m_{e_{mil}}$ and $m_{e_{path}}$) together with interquartile ranges ($IQR_{e_{mil}}$ and $IQR_{e_{path}}$) are presented.

| | MIL prediction | | | | Pathologist's estimate | | | |
|---|---|---|---|---|---|---|---|---|
| | $\mu_{e_{mil}}$ | $\sigma_{e_{mil}}$ | $m_{e_{mil}}$ | $IQR_{e_{mil}}$ | $\mu_{e_{path}}$ | $\sigma_{e_{path}}$ | $m_{e_{path}}$ | $IQR_{e_{path}}$ |
| training | 0.069 | 0.073 | 0.046 | 0.019 - 0.100 | 0.276 | 0.155 | 0.270 | 0.150 - 0.380 |
| validation | 0.123 | 0.122 | 0.093 | 0.040 - 0.159 | 0.256 | 0.140 | 0.250 | 0.135 - 0.350 |
| test | 0.132 | 0.109 | 0.112 | 0.060 - 0.175 | 0.280 | 0.151 | 0.275 | 0.170 - 0.395 |

Table 27: **LUSC cohort.** Absolute errors between genomic tumor purity values and MIL predictions ($e_{mil}$) and genomic tumor purity values and pathologists' percent tumor nuclei estimates ($e_{path}$) in the training, validation, and test sets are calculated using only the tumor samples. Mean absolute errors ($\mu_{e_{mil}}$ and $\mu_{e_{path}}$) together with standard deviations ($\sigma_{e_{mil}}$ and $\sigma_{e_{path}}$) and median absolute errors ($m_{e_{mil}}$ and $m_{e_{path}}$) together with interquartile ranges ($IQR_{e_{mil}}$ and $IQR_{e_{path}}$) are presented.

| | MIL prediction | | | | Pathologist's estimate | | | |
|---|---|---|---|---|---|---|---|---|
| | $\mu_{e_{mil}}$ | $\sigma_{e_{mil}}$ | $m_{e_{mil}}$ | $IQR_{e_{mil}}$ | $\mu_{e_{path}}$ | $\sigma_{e_{path}}$ | $m_{e_{path}}$ | $IQR_{e_{path}}$ |
| training | 0.093 | 0.080 | 0.078 | 0.038 - 0.134 | 0.254 | 0.151 | 0.250 | 0.130 - 0.370 |
| validation | 0.134 | 0.103 | 0.111 | 0.064 - 0.183 | 0.263 | 0.147 | 0.260 | 0.140 - 0.370 |
| test | 0.148 | 0.122 | 0.125 | 0.054 - 0.196 | 0.266 | 0.150 | 0.250 | 0.140 - 0.375 |

Table 28: **OV cohort.** Absolute errors between genomic tumor purity values and MIL predictions ($e_{mil}$) and genomic tumor purity values and pathologists' percent tumor nuclei estimates ($e_{path}$) in the training, validation, and test sets are calculated using only the tumor samples. Mean absolute errors ($\mu_{e_{mil}}$ and $\mu_{e_{path}}$) together with standard deviations ($\sigma_{e_{mil}}$ and $\sigma_{e_{path}}$) and median absolute errors ($m_{e_{mil}}$ and $m_{e_{path}}$) together with interquartile ranges ($IQR_{e_{mil}}$ and $IQR_{e_{path}}$) are presented.

| | MIL prediction | | | | Pathologist's estimate | | | |
|---|---|---|---|---|---|---|---|---|
| | $\mu_{e_{mil}}$ | $\sigma_{e_{mil}}$ | $m_{e_{mil}}$ | $IQR_{e_{mil}}$ | $\mu_{e_{path}}$ | $\sigma_{e_{path}}$ | $m_{e_{path}}$ | $IQR_{e_{path}}$ |
| training | 0.042 | 0.038 | 0.033 | 0.018 - 0.056 | 0.131 | 0.119 | 0.100 | 0.040 - 0.190 |
| validation | 0.101 | 0.093 | 0.076 | 0.040 - 0.128 | 0.145 | 0.134 | 0.100 | 0.055 - 0.190 |
| test | 0.105 | 0.091 | 0.086 | 0.043 - 0.127 | 0.136 | 0.126 | 0.110 | 0.030 - 0.190 |

Table 29: **PRAD cohort.** Absolute errors between genomic tumor purity values and MIL predictions ($e_{mil}$) and genomic tumor purity values and pathologists' percent tumor nuclei estimates ($e_{path}$) in the training, validation, and test sets are calculated using only the tumor samples. Mean absolute errors ($\mu_{e_{mil}}$ and $\mu_{e_{path}}$) together with standard deviations ($\sigma_{e_{mil}}$ and $\sigma_{e_{path}}$) and median absolute errors ($m_{e_{mil}}$ and $m_{e_{path}}$) together with interquartile ranges ($IQR_{e_{mil}}$ and $IQR_{e_{path}}$) are presented.

| | MIL prediction | | | | Pathologist's estimate | | | |
|---|---|---|---|---|---|---|---|---|
| | $\mu_{e_{mil}}$ | $\sigma_{e_{mil}}$ | $m_{e_{mil}}$ | $IQR_{e_{mil}}$ | $\mu_{e_{path}}$ | $\sigma_{e_{path}}$ | $m_{e_{path}}$ | $IQR_{e_{path}}$ |
| training | 0.166 | 0.141 | 0.128 | 0.065 - 0.235 | 0.203 | 0.147 | 0.170 | 0.080 - 0.310 |
| validation | 0.179 | 0.137 | 0.161 | 0.077 - 0.254 | 0.214 | 0.157 | 0.180 | 0.085 - 0.340 |
| test | 0.173 | 0.154 | 0.130 | 0.068 - 0.240 | 0.204 | 0.141 | 0.180 | 0.090 - 0.285 |

Table 30: **THCA cohort.** Absolute errors between genomic tumor purity values and MIL predictions ($e_{mil}$) and genomic tumor purity values and pathologists' percent tumor nuclei estimates ($e_{path}$) in the training, validation, and test sets are calculated using only the tumor samples. Mean absolute errors ($\mu_{e_{mil}}$ and $\mu_{e_{path}}$) together with standard deviations ($\sigma_{e_{mil}}$ and $\sigma_{e_{path}}$) and median absolute errors ($m_{e_{mil}}$ and $m_{e_{path}}$) together with interquartile ranges ($IQR_{e_{mil}}$ and $IQR_{e_{path}}$) are presented.

| | MIL prediction | | | | Pathologist's estimate | | | |
|---|---|---|---|---|---|---|---|---|
| | $\mu_{e_{mil}}$ | $\sigma_{e_{mil}}$ | $m_{e_{mil}}$ | $IQR_{e_{mil}}$ | $\mu_{e_{path}}$ | $\sigma_{e_{path}}$ | $m_{e_{path}}$ | $IQR_{e_{path}}$ |
| training | 0.082 | 0.083 | 0.058 | 0.022 - 0.119 | 0.213 | 0.147 | 0.180 | 0.100 - 0.300 |
| validation | 0.133 | 0.095 | 0.118 | 0.064 - 0.188 | 0.205 | 0.148 | 0.170 | 0.110 - 0.290 |
| test | 0.171 | 0.126 | 0.145 | 0.070 - 0.258 | 0.205 | 0.140 | 0.180 | 0.090 - 0.310 |

Table 31: **UCEC cohort.** Absolute errors between genomic tumor purity values and MIL predictions ($e_{mil}$) and genomic tumor purity values and pathologists' percent tumor nuclei estimates ($e_{path}$) in the training, validation, and test sets are calculated using only the tumor samples. Mean absolute errors ($\mu_{e_{mil}}$ and $\mu_{e_{path}}$) together with standard deviations ($\sigma_{e_{mil}}$ and $\sigma_{e_{path}}$) and median absolute errors ($m_{e_{mil}}$ and $m_{e_{path}}$) together with interquartile ranges ($IQR_{e_{mil}}$ and $IQR_{e_{path}}$) are presented.

| | MIL prediction | | | | Pathologist's estimate | | | |
|---|---|---|---|---|---|---|---|---|
| | $\mu_{e_{mil}}$ | $\sigma_{e_{mil}}$ | $m_{e_{mil}}$ | $IQR_{e_{mil}}$ | $\mu_{e_{path}}$ | $\sigma_{e_{path}}$ | $m_{e_{path}}$ | $IQR_{e_{path}}$ |
| training | 0.073 | 0.092 | 0.049 | 0.019 - 0.090 | 0.140 | 0.133 | 0.100 | 0.040 - 0.190 |
| validation | 0.114 | 0.109 | 0.087 | 0.047 - 0.134 | 0.134 | 0.137 | 0.090 | 0.040 - 0.180 |
| test | 0.109 | 0.120 | 0.072 | 0.027 - 0.142 | 0.132 | 0.124 | 0.100 | 0.040 - 0.170 |

Table 32: **Comparison of methods based on absolute errors in the test sets of different cohorts.** Absolute errors between genomic tumor purity values and MIL predictions ($e_{mil}$) and genomic tumor purity values and pathologists' percent tumor nuclei estimates ($e_{path}$) in the test sets of different cohorts are calculated for only the tumor samples. Then, they are compared using the Wilcoxon signed-rank test [3]. Mean absolute errors ($\mu_{e_{mil}}$ and $\mu_{e_{path}}$) together with standard deviations ($\sigma_{e_{mil}}$ and $\sigma_{e_{path}}$), median absolute errors ($m_{e_{mil}}$ and $m_{e_{path}}$) together with interquartile ranges ($IQR_{e_{mil}}$ and $IQR_{e_{path}}$), and calculated p-values in the statistical tests ($P_{comp}$) are presented. The best methods are highlighted in bold.

| | MIL prediction | | | | Pathologist's estimate | | | | Comparison |
|---|---|---|---|---|---|---|---|---|---|
| | $\mu_{e_{mil}}$ | $\sigma_{e_{mil}}$ | $m_{e_{mil}}$ | $IQR_{e_{mil}}$ | $\mu_{e_{path}}$ | $\sigma_{e_{path}}$ | $m_{e_{path}}$ | $IQR_{e_{path}}$ | $P_{comp}$ |
| BRCA | **0.116** | **0.097** | **0.104** | **0.043 - 0.159** | 0.220 | 0.147 | 0.200 | 0.105 - 0.310 | **2.5e-13** |
| GBM | **0.113** | **0.106** | **0.074** | **0.046 - 0.142** | 0.195 | 0.158 | 0.145 | 0.080 - 0.260 | **2.1e-07** |
| LGG | 0.136 | 0.119 | 0.105 | 0.052 - 0.188 | 0.152 | 0.122 | 0.130 | 0.060 - 0.200 | 5.4e-02 |
| LUAD | **0.132** | **0.109** | **0.112** | **0.060 - 0.175** | 0.280 | 0.151 | 0.275 | 0.170 - 0.395 | **3.9e-09** |
| LUSC | **0.148** | **0.122** | **0.125** | **0.054 - 0.196** | 0.266 | 0.150 | 0.250 | 0.140 - 0.375 | **5.8e-06** |
| OV | **0.105** | **0.091** | **0.086** | **0.043 - 0.127** | 0.136 | 0.126 | 0.110 | 0.030 - 0.190 | **1.6e-02** |
| PRAD | **0.173** | **0.154** | **0.130** | **0.068 - 0.240** | 0.204 | 0.141 | 0.180 | 0.090 - 0.285 | **1.4e-02** |
| UCEC | **0.109** | **0.120** | **0.072** | **0.027 - 0.142** | 0.132 | 0.124 | 0.100 | 0.040 - 0.170 | **1.4e-02** |

#### 2.3 Tumor Purity Prediction Differences Between Slides of The Same Sample

For a tumor sample with two slides, let $\hat{p}_{sld1}$ and $\hat{p}_{sld2}$ be tumor purity predictions obtained from the trained MIL model for the slides of the tumor sample. Then, the absolute difference between the predictions is calculated as $d_{abs} = |\hat{p}_{sld1} - \hat{p}_{sld2}|$. Histograms of absolute differences in training, validation, and test sets are presented for: BRCA cohort in Figure 21, GBM cohort in Figure 22, LGG cohort in Figure 23, LUAD cohort in Figure 24, LUSC cohort in Figure 25, OV cohort in Figure 26, PRAD cohort in Figure 27, UCEC cohort in Figure 28.

Moreover, for each set, the number of tumor samples with two slides (n), the mean absolute difference ($\mu_{d_{abs}}$), the standard deviation of absolute difference ($\sigma_{d_{abs}}$), the median absolute difference ($m_{d_{abs}}$), and the interquartile range ($IQR_{d_{abs}}$) are given for: BRCA cohort in Table 33, GBM cohort in Table 34, LGG cohort in Table 35, LUAD cohort in Table 36, LUSC cohort in Table 37, OV cohort in Table 38, PRAD cohort in Table 39, UCEC cohort in Table 40.

We summarized the number of tumor samples with two slides (n), the mean absolute difference ($\mu_{d_{abs}}$), the standard deviation of absolute difference ($\sigma_{d_{abs}}$), the median absolute difference ($m_{d_{abs}}$), and the interquartile range ($IQR_{d_{abs}}$) for the test set of each cohort in Table 41.

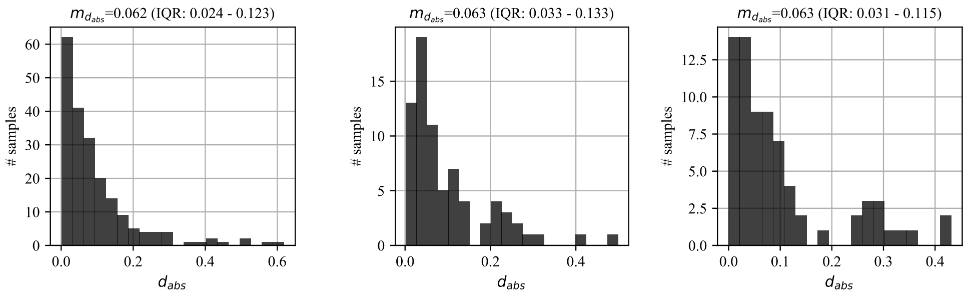


(a) Training (b) Validation (c) Test

Figure 21: **Histograms of the absolute difference between slides of the same sample - BRCA cohort.** In the (a) training, (b) validation, and (c) test sets, for a tumor sample with two slides, absolute difference ($d_{abs}$) between the slides' tumor purity predictions is calculated. Then, a histogram of absolute differences is presented. Note that median absolute difference ($m_{d_{abs}}$) and interquartile range ($IQR_{d_{abs}}$) are also given in the title of each chart.

Table 33: **Statistics of the absolute difference between slides of the same sample - BRCA cohort.** In the training, validation, and test sets of the cohort, for a tumor sample with two slides, the absolute difference ($d_{abs}$) between the slides' tumor purity predictions is calculated. Then, the number of tumor samples with two slides (n), the mean absolute difference ($\mu_{d_{abs}}$), the standard deviation of absolute difference ($\sigma_{d_{abs}}$), the median absolute difference ($m_{d_{abs}}$), and the interquartile range ($IQR_{d_{abs}}$) are presented.

| | n | $\mu_{d_{abs}}$ | $\sigma_{d_{abs}}$ | $m_{d_{abs}}$ | $IQR_{d_{abs}}$ |
|---|---|---|---|---|---|
| training | 204 | 0.095 | 0.107 | 0.062 | 0.024 - 0.123 |
| validation | 74 | 0.099 | 0.099 | 0.063 | 0.033 - 0.133 |
| test | 73 | 0.101 | 0.106 | 0.063 | 0.031 - 0.115 |

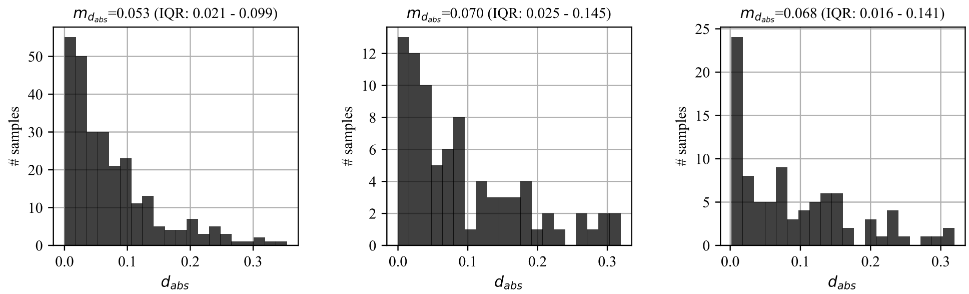


(a) Training (b) Validation (c) Test

Figure 22: **Histograms of the absolute difference between slides of the same sample - GBM cohort.** In the (a) training, (b) validation, and (c) test sets, for a tumor sample with two slides, absolute difference ($d_{abs}$) between the slides' tumor purity predictions is calculated. Then, a histogram of absolute differences is presented. Note that median absolute difference ($m_{d_{abs}}$) and interquartile range ($IQR_{d_{abs}}$) are also given in the title of each chart.

Table 34: **Statistics of the absolute difference between slides of the same sample - GBM cohort.** In the training, validation, and test sets of the cohort, for a tumor sample with two slides, the absolute difference ($d_{abs}$) between the slides' tumor purity predictions is calculated. Then, the number of tumor samples with two slides (n), the mean absolute difference ($\mu_{d_{abs}}$), the standard deviation of absolute difference ($\sigma_{d_{abs}}$), the median absolute difference ($m_{d_{abs}}$), and the interquartile range ($IQR_{d_{abs}}$) are presented.

| | n | $\mu_{d_{abs}}$ | $\sigma_{d_{abs}}$ | $m_{d_{abs}}$ | $IQR_{d_{abs}}$ |
|---|---|---|---|---|---|
| training | 270 | 0.074 | 0.070 | 0.053 | 0.021 - 0.099 |
| validation | 83 | 0.093 | 0.085 | 0.070 | 0.025 - 0.145 |
| test | 90 | 0.090 | 0.083 | 0.068 | 0.016 - 0.141 |

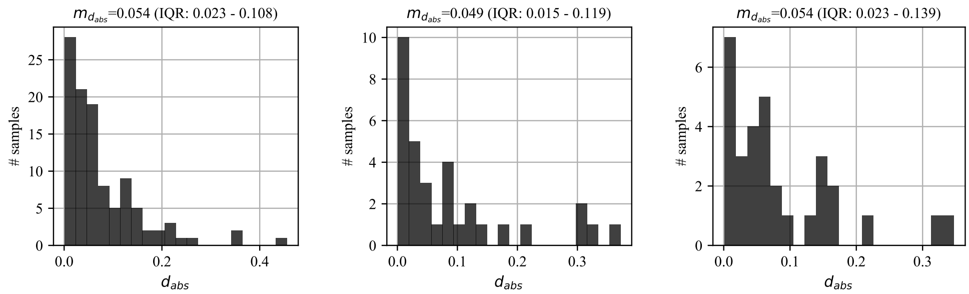


(a) Training (b) Validation (c) Test

Figure 23: **Histograms of the absolute difference between slides of the same sample - LGG cohort.** In the (a) training, (b) validation, and (c) test sets, for a tumor sample with two slides, absolute difference ($d_{abs}$) between the slides' tumor purity predictions is calculated. Then, a histogram of absolute differences is presented. Note that median absolute difference ($m_{d_{abs}}$) and interquartile range ($IQR_{d_{abs}}$) are also given in the title of each chart.

Table 35: **Statistics of the absolute difference between slides of the same sample - LGG cohort.** In the training, validation, and test sets of the cohort, for a tumor sample with two slides, the absolute difference ($d_{abs}$) between the slides' tumor purity predictions is calculated. Then, the number of tumor samples with two slides (n), the mean absolute difference ($\mu_{d_{abs}}$), the standard deviation of absolute difference ($\sigma_{d_{abs}}$), the median absolute difference ($m_{d_{abs}}$), and the interquartile range ($IQR_{d_{abs}}$) are presented.

| | n | $\mu_{d_{abs}}$ | $\sigma_{d_{abs}}$ | $m_{d_{abs}}$ | $IQR_{d_{abs}}$ |
|---|---|---|---|---|---|
| training | 107 | 0.077 | 0.079 | 0.054 | 0.023 - 0.108 |
| validation | 33 | 0.091 | 0.104 | 0.049 | 0.015 - 0.119 |
| test | 31 | 0.086 | 0.089 | 0.054 | 0.023 - 0.139 |

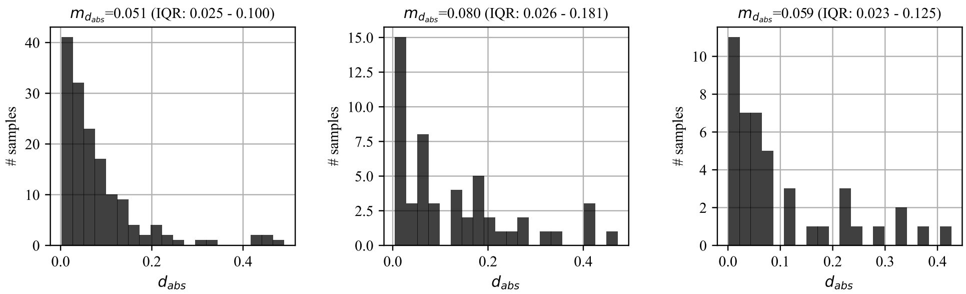


(a) Training (b) Validation (c) Test

Figure 24: **Histograms of the absolute difference between slides of the same sample - LUAD cohort.** In the (a) training, (b) validation, and (c) test sets, for a tumor sample with two slides, absolute difference ($d_{abs}$) between the slides' tumor purity predictions is calculated. Then, a histogram of absolute differences is presented. Note that median absolute difference ($m_{d_{abs}}$) and interquartile range ($IQR_{d_{abs}}$) are also given in the title of each chart.

Table 36: **Statistics of the absolute difference between slides of the same sample - LUAD cohort.** In the training, validation, and test sets of the cohort, for a tumor sample with two slides, the absolute difference ($d_{abs}$) between the slides' tumor purity predictions is calculated. Then, the number of tumor samples with two slides (n), the mean absolute difference ($\mu_{d_{abs}}$), the standard deviation of absolute difference ($\sigma_{d_{abs}}$), the median absolute difference ($m_{d_{abs}}$), and the interquartile range ($IQR_{d_{abs}}$) are presented.

| | n | $\mu_{d_{abs}}$ | $\sigma_{d_{abs}}$ | $m_{d_{abs}}$ | $IQR_{d_{abs}}$ |
|---|---|---|---|---|---|
| training | 152 | 0.082 | 0.092 | 0.051 | 0.025 - 0.100 |
| validation | 52 | 0.127 | 0.123 | 0.080 | 0.026 - 0.181 |
| test | 44 | 0.100 | 0.110 | 0.059 | 0.023 - 0.125 |

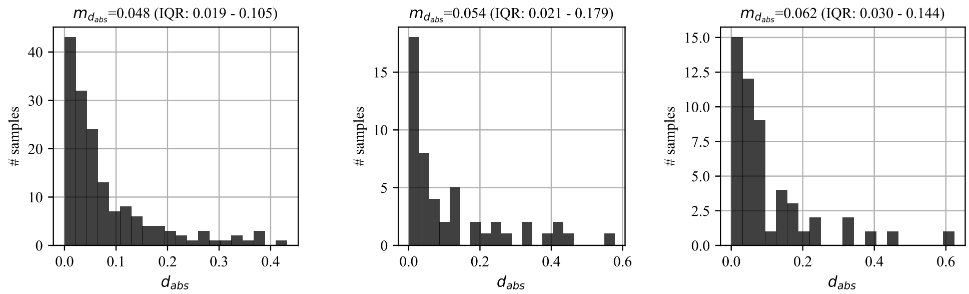


(a) Training (b) Validation (c) Test

Figure 25: **Histograms of the absolute difference between slides of the same sample - LUSC cohort.** In the (a) training, (b) validation, and (c) test sets, for a tumor sample with two slides, absolute difference ($d_{abs}$) between the slides' tumor purity predictions is calculated. Then, a histogram of absolute differences is presented. Note that median absolute difference ($m_{d_{abs}}$) and interquartile range ($IQR_{d_{abs}}$) are also given in the title of each chart.

Table 37: **Statistics of the absolute difference between slides of the same sample - LUSC cohort.** In the training, validation, and test sets of the cohort, for a tumor sample with two slides, the absolute difference ($d_{abs}$) between the slides' tumor purity predictions is calculated. Then, the number of tumor samples with two slides (n), the mean absolute difference ($\mu_{d_{abs}}$), the standard deviation of absolute difference ($\sigma_{d_{abs}}$), the median absolute difference ($m_{d_{abs}}$), and the interquartile range ($IQR_{d_{abs}}$) are presented.

| | n | $\mu_{d_{abs}}$ | $\sigma_{d_{abs}}$ | $m_{d_{abs}}$ | $IQR_{d_{abs}}$ |
|---|---|---|---|---|---|
| training | 159 | 0.080 | 0.092 | 0.048 | 0.019 - 0.105 |
| validation | 50 | 0.120 | 0.142 | 0.054 | 0.021 - 0.179 |
| test | 52 | 0.106 | 0.123 | 0.062 | 0.030 - 0.144 |

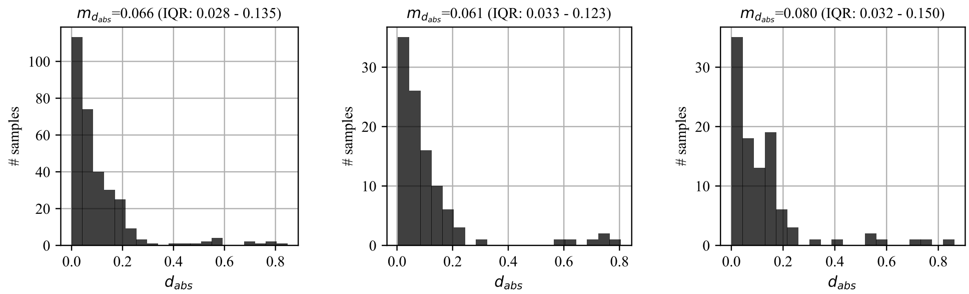


(a) Training (b) Validation (c) Test

Figure 26: **Histograms of the absolute difference between slides of the same sample - OV cohort.** In the (a) training, (b) validation, and (c) test sets, for a tumor sample with two slides, absolute difference ($d_{abs}$) between the slides' tumor purity predictions is calculated. Then, a histogram of absolute differences is presented. Note that median absolute difference ($m_{d_{abs}}$) and interquartile range ($IQR_{d_{abs}}$) are also given in the title of each chart.

Table 38: **Statistics of the absolute difference between slides of the same sample - OV cohort.** In the training, validation, and test sets of the cohort, for a tumor sample with two slides, the absolute difference ($d_{abs}$) between the slides' tumor purity predictions is calculated. Then, the number of tumor samples with two slides (n), the mean absolute difference ($\mu_{d_{abs}}$), the standard deviation of absolute difference ($\sigma_{d_{abs}}$), the median absolute difference ($m_{d_{abs}}$), and the interquartile range ($IQR_{d_{abs}}$) are presented.

| | n | $\mu_{d_{abs}}$ | $\sigma_{d_{abs}}$ | $m_{d_{abs}}$ | $IQR_{d_{abs}}$ |
|---|---|---|---|---|---|
| training | 310 | 0.106 | 0.135 | 0.066 | 0.028 - 0.135 |
| validation | 103 | 0.113 | 0.157 | 0.061 | 0.031 - 0.123 |
| test | 102 | 0.125 | 0.156 | 0.080 | 0.032 - 0.150 |

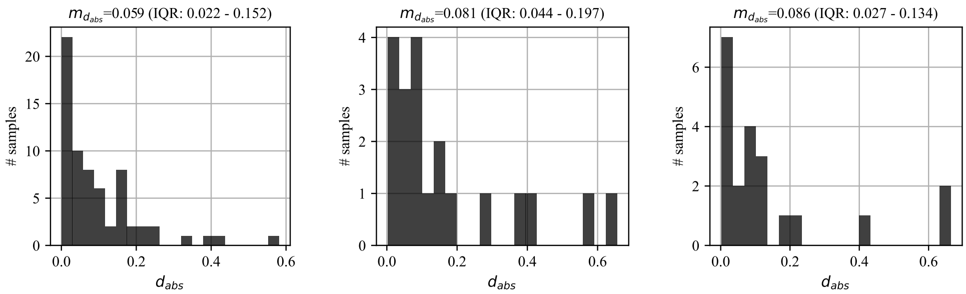


(a) Training (b) Validation (c) Test

Figure 27: **Histograms of the absolute difference between slides of the same sample - PRAD cohort.** In the (a) training, (b) validation, and (c) test sets, for a tumor sample with two slides, absolute difference ($d_{abs}$) between the slides' tumor purity predictions is calculated. Then, a histogram of absolute differences is presented. Note that median absolute difference ($m_{d_{abs}}$) and interquartile range ($IQR_{d_{abs}}$) are also given in the title of each chart.

Table 39: **Statistics of the absolute difference between slides of the same sample - PRAD cohort.** In the training, validation, and test sets of the cohort, for a tumor sample with two slides, the absolute difference ($d_{abs}$) between the slides' tumor purity predictions is calculated. Then, the number of tumor samples with two slides (n), the mean absolute difference ($\mu_{d_{abs}}$), the standard deviation of absolute difference ($\sigma_{d_{abs}}$), the median absolute difference ($m_{d_{abs}}$), and the interquartile range ($IQR_{d_{abs}}$) are presented.

| | n | $\mu_{d_{abs}}$ | $\sigma_{d_{abs}}$ | $m_{d_{abs}}$ | $IQR_{d_{abs}}$ |
|---|---|---|---|---|---|
| training | 66 | 0.098 | 0.110 | 0.059 | 0.022 - 0.152 |
| validation | 20 | 0.174 | 0.186 | 0.081 | 0.044 - 0.197 |
| test | 21 | 0.144 | 0.189 | 0.086 | 0.027 - 0.134 |

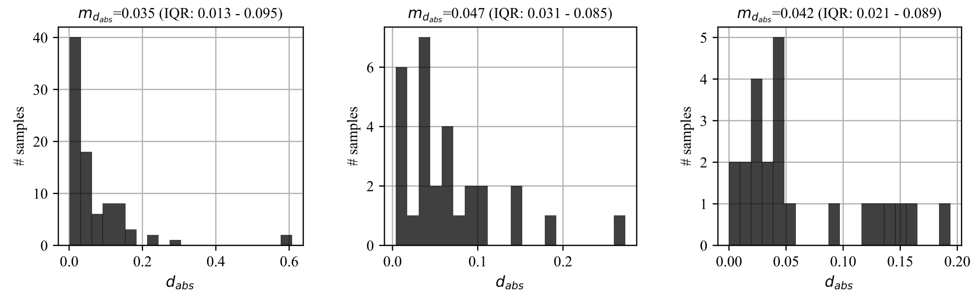


(a) Training (b) Validation (c) Test

Figure 28: **Histograms of the absolute difference between slides of the same sample - UCEC cohort.** In the (a) training, (b) validation, and (c) test sets, for a tumor sample with two slides, absolute difference ($d_{abs}$) between the slides' tumor purity predictions is calculated. Then, a histogram of absolute differences is presented. Note that median absolute difference ($m_{d_{abs}}$) and interquartile range ($IQR_{d_{abs}}$) are also given in the title of each chart.

Table 40: **Statistics of the absolute difference between slides of the same sample - UCEC cohort.** In the training, validation, and test sets of the cohort, for a tumor sample with two slides, the absolute difference ($d_{abs}$) between the slides' tumor purity predictions is calculated. Then, the number of tumor samples with two slides (n), the mean absolute difference ($\mu_{d_{abs}}$), the standard deviation of absolute difference ($\sigma_{d_{abs}}$), the median absolute difference ($m_{d_{abs}}$), and the interquartile range ($IQR_{d_{abs}}$) are presented.

| | n | $\mu_{d_{abs}}$ | $\sigma_{d_{abs}}$ | $m_{d_{abs}}$ | $IQR_{d_{abs}}$ |
|---|---|---|---|---|---|
| training | 88 | 0.069 | 0.100 | 0.035 | 0.013 - 0.095 |
| validation | 29 | 0.066 | 0.060 | 0.047 | 0.031 - 0.085 |
| test | 23 | 0.063 | 0.056 | 0.042 | 0.021 - 0.089 |

Table 41: **Statistics of the absolute difference between the predictions of a tumor sample's top and bottom slides.** In the test set of each cohort, for a tumor sample with two slides, the absolute difference ($d_{abs}$) between the tumor purity predictions of the slides is calculated. Then, the number of tumor samples with two slides (n), the mean absolute difference ($\mu_{d_{abs}}$), the standard deviation of the absolute difference ($\sigma_{d_{abs}}$), the median absolute difference ($m_{d_{abs}}$), and the interquartile range ($IQR_{d_{abs}}$) are presented.

| | n | $\mu_{d_{abs}}$ | $\sigma_{d_{abs}}$ | $m_{d_{abs}}$ | $IQR_{d_{abs}}$ |
|---|---|---|---|---|---|
| BRCA | 73 | 0.101 | 0.106 | 0.063 | 0.031 - 0.115 |
| GBM | 90 | 0.090 | 0.083 | 0.068 | 0.016 - 0.141 |
| LGG | 31 | 0.086 | 0.089 | 0.054 | 0.023 - 0.139 |
| LUAD | 44 | 0.100 | 0.110 | 0.059 | 0.023 - 0.125 |
| LUSC | 52 | 0.106 | 0.123 | 0.062 | 0.030 - 0.144 |
| OV | 102 | 0.125 | 0.156 | 0.080 | 0.032 - 0.150 |
| PRAD | 21 | 0.144 | 0.189 | 0.086 | 0.027 - 0.134 |
| UCEC | 23 | 0.063 | 0.056 | 0.042 | 0.021 - 0.089 |

#### 2.4 Predicting sample-level tumor purity using two slides together is better than using one of the slides

For a tumor sample with two slides, let $p_{smpl}$ be the sample's genomic tumor purity obtained from ABSOLUTE; $\hat{p}_{smpl}$ be sample-level tumor purity prediction obtained from the trained MIL model using both of the slides together; $\hat{p}_{sld1}$ and $\hat{p}_{sld2}$ be tumor purity predictions obtained from the trained MIL model for individual slides. We compare the absolute error of sample-level prediction ($e_{smpl} = |\hat{p}_{smpl} - p_{smpl}|$) and expected value of absolute errors of slide-level predictions ($e_{sld} = 0.5 * (|\hat{p}_{sld1} - p_{smpl}| + |\hat{p}_{sld2} - p_{smpl}|)$). We use the Wilcoxon signed-rank test [3] on the difference of $e_{smpl} - e_{sld}$.

The results of our analyses in the training, validation, and test sets are presented for: BRCA cohort in Table 42, GBM cohort in Table 43, LGG cohort in Table 44, LUAD cohort in Table 45, LUSC cohort in Table 46, OV cohort in Table 47, PRAD cohort in Table 48, UCEC cohort in Table 49.

We also summarize the results of our analyses in the test sets of different cohorts in Table 50.

Table 42: **Absolute error analysis: using two slides together vs. using one of the slides to predict sample-level tumor purity - BRCA cohort.** For a tumor sample with two slides, absolute errors between genomic tumor purity values and sample-level MIL predictions ($e_{smpl}$) and genomic tumor purity values and slide-level MIL predictions ($e_{sld}$) in the training, validation, and test sets are calculated. Mean absolute errors ($\mu_{e_{smpl}}$ and $\mu_{e_{sld}}$) together with standard deviations ($\sigma_{e_{smpl}}$ and $\sigma_{e_{sld}}$) and median absolute errors ($m_{e_{smpl}}$ and $m_{e_{sld}}$) together with interquartile ranges ($IQR_{e_{smpl}}$ and $IQR_{e_{sld}}$) are presented.

| | sample level | | | | slide level | | | |
|---|---|---|---|---|---|---|---|---|
| | $\mu_{e_{smpl}}$ | $\sigma_{e_{smpl}}$ | $m_{e_{smpl}}$ | $IQR_{e_{smpl}}$ | $\mu_{e_{sld}}$ | $\sigma_{e_{sld}}$ | $m_{e_{sld}}$ | $IQR_{e_{sld}}$ |
| training | 0.089 | 0.066 | 0.076 | 0.036 - 0.125 | 0.100 | 0.075 | 0.082 | 0.043 - 0.138 |
| validation | 0.127 | 0.132 | 0.090 | 0.035 - 0.187 | 0.134 | 0.133 | 0.088 | 0.031 - 0.202 |
| test | 0.114 | 0.082 | 0.092 | 0.043 - 0.166 | 0.113 | 0.078 | 0.101 | 0.048 - 0.166 |

Table 43: **Absolute error analysis: using two slides together vs. using one of the slides to predict sample-level tumor purity - GBM cohort.** For a tumor sample with two slides, absolute errors between genomic tumor purity values and sample-level MIL predictions ($e_{smpl}$) and genomic tumor purity values and slide-level MIL predictions ($e_{sld}$) in the training, validation, and test sets are calculated. Mean absolute errors ($\mu_{e_{smpl}}$ and $\mu_{e_{sld}}$) together with standard deviations ($\sigma_{e_{smpl}}$ and $\sigma_{e_{sld}}$) and median absolute errors ($m_{e_{smpl}}$ and $m_{e_{sld}}$) together with interquartile ranges ($IQR_{e_{smpl}}$ and $IQR_{e_{sld}}$) are presented.

| | sample level | | | | slide level | | | |
|---|---|---|---|---|---|---|---|---|
| | $\mu_{e_{smpl}}$ | $\sigma_{e_{smpl}}$ | $m_{e_{smpl}}$ | $IQR_{e_{smpl}}$ | $\mu_{e_{sld}}$ | $\sigma_{e_{sld}}$ | $m_{e_{sld}}$ | $IQR_{e_{sld}}$ |
| training | 0.054 | 0.059 | 0.032 | 0.017 - 0.066 | 0.089 | 0.070 | 0.070 | 0.036 - 0.120 |
| validation | 0.107 | 0.095 | 0.080 | 0.023 - 0.164 | 0.115 | 0.084 | 0.090 | 0.045 - 0.165 |
| test | 0.115 | 0.107 | 0.076 | 0.046 - 0.145 | 0.118 | 0.096 | 0.089 | 0.062 - 0.161 |

Table 44: **Absolute error analysis: using two slides together vs. using one of the slides to predict sample-level tumor purity - LGG cohort.** For a tumor sample with two slides, absolute errors between genomic tumor purity values and sample-level MIL predictions ($e_{smpl}$) and genomic tumor purity values and slide-level MIL predictions ($e_{sld}$) in the training, validation, and test sets are calculated. Mean absolute errors ($\mu_{e_{smpl}}$ and $\mu_{e_{sld}}$) together with standard deviations ($\sigma_{e_{smpl}}$ and $\sigma_{e_{sld}}$) and median absolute errors ($m_{e_{smpl}}$ and $m_{e_{sld}}$) together with interquartile ranges ($IQR_{e_{smpl}}$ and $IQR_{e_{sld}}$) are presented.

| | sample level | | | | slide level | | | |
|---|---|---|---|---|---|---|---|---|
| | $\mu_{e_{smpl}}$ | $\sigma_{e_{smpl}}$ | $m_{e_{smpl}}$ | $IQR_{e_{smpl}}$ | $\mu_{e_{sld}}$ | $\sigma_{e_{sld}}$ | $m_{e_{sld}}$ | $IQR_{e_{sld}}$ |
| training | 0.079 | 0.080 | 0.053 | 0.028 - 0.109 | 0.118 | 0.114 | 0.079 | 0.043 - 0.143 |
| validation | 0.133 | 0.137 | 0.096 | 0.027 - 0.201 | 0.149 | 0.133 | 0.123 | 0.056 - 0.164 |
| test | 0.178 | 0.149 | 0.146 | 0.100 - 0.218 | 0.168 | 0.152 | 0.106 | 0.067 - 0.198 |

Table 45: **Absolute error analysis: using two slides together vs. using one of the slides to predict sample-level tumor purity - LUAD cohort.** For a tumor sample with two slides, absolute errors between genomic tumor purity values and sample-level MIL predictions ($e_{smpl}$) and genomic tumor purity values and slide-level MIL predictions ($e_{sld}$) in the training, validation, and test sets are calculated. Mean absolute errors ($\mu_{e_{smpl}}$ and $\mu_{e_{sld}}$) together with standard deviations ($\sigma_{e_{smpl}}$ and $\sigma_{e_{sld}}$) and median absolute errors ($m_{e_{smpl}}$ and $m_{e_{sld}}$) together with interquartile ranges ($IQR_{e_{smpl}}$ and $IQR_{e_{sld}}$) are presented.

| | sample level | | | | slide level | | | |
|---|---|---|---|---|---|---|---|---|
| | $\mu_{e_{smpl}}$ | $\sigma_{e_{smpl}}$ | $m_{e_{smpl}}$ | $IQR_{e_{smpl}}$ | $\mu_{e_{sld}}$ | $\sigma_{e_{sld}}$ | $m_{e_{sld}}$ | $IQR_{e_{sld}}$ |
| training | 0.064 | 0.058 | 0.047 | 0.019 - 0.089 | 0.084 | 0.066 | 0.067 | 0.035 - 0.108 |
| validation | 0.121 | 0.107 | 0.096 | 0.047 - 0.173 | 0.146 | 0.110 | 0.129 | 0.060 - 0.207 |
| test | 0.118 | 0.102 | 0.084 | 0.050 - 0.168 | 0.138 | 0.102 | 0.121 | 0.067 - 0.181 |

Table 46: **Absolute error analysis: using two slides together vs. using one of the slides to predict sample-level tumor purity - LUSC cohort.** For a tumor sample with two slides, absolute errors between genomic tumor purity values and sample-level MIL predictions ($e_{smpl}$) and genomic tumor purity values and slide-level MIL predictions ($e_{sld}$) in the training, validation, and test sets are calculated. Mean absolute errors ($\mu_{e_{smpl}}$ and $\mu_{e_{sld}}$) together with standard deviations ($\sigma_{e_{smpl}}$ and $\sigma_{e_{sld}}$) and median absolute errors ($m_{e_{smpl}}$ and $m_{e_{sld}}$) together with interquartile ranges ($IQR_{e_{smpl}}$ and $IQR_{e_{sld}}$) are presented.

| | sample level | | | | slide level | | | |
|---|---|---|---|---|---|---|---|---|
| | $\mu_{e_{smpl}}$ | $\sigma_{e_{smpl}}$ | $m_{e_{smpl}}$ | $IQR_{e_{smpl}}$ | $\mu_{e_{sld}}$ | $\sigma_{e_{sld}}$ | $m_{e_{sld}}$ | $IQR_{e_{sld}}$ |
| training | 0.090 | 0.069 | 0.071 | 0.038 - 0.132 | 0.128 | 0.097 | 0.107 | 0.056 - 0.181 |
| validation | 0.137 | 0.107 | 0.111 | 0.061 - 0.199 | 0.157 | 0.102 | 0.138 | 0.071 - 0.229 |
| test | 0.124 | 0.092 | 0.109 | 0.039 - 0.168 | 0.150 | 0.096 | 0.143 | 0.085 - 0.201 |

Table 47: **Absolute error analysis: using two slides together vs. using one of the slides to predict sample-level tumor purity - OV cohort.** For a tumor sample with two slides, absolute errors between genomic tumor purity values and sample-level MIL predictions ($e_{smpl}$) and genomic tumor purity values and slide-level MIL predictions ($e_{sld}$) in the training, validation, and test sets are calculated. Mean absolute errors ($\mu_{e_{smpl}}$ and $\mu_{e_{sld}}$) together with standard deviations ($\sigma_{e_{smpl}}$ and $\sigma_{e_{sld}}$) and median absolute errors ($m_{e_{smpl}}$ and $m_{e_{sld}}$) together with interquartile ranges ($IQR_{e_{smpl}}$ and $IQR_{e_{sld}}$) are presented.

| | sample level | | | | slide level | | | |
|---|---|---|---|---|---|---|---|---|
| | $\mu_{e_{smpl}}$ | $\sigma_{e_{smpl}}$ | $m_{e_{smpl}}$ | $IQR_{e_{smpl}}$ | $\mu_{e_{sld}}$ | $\sigma_{e_{sld}}$ | $m_{e_{sld}}$ | $IQR_{e_{sld}}$ |
| training | 0.042 | 0.038 | 0.033 | 0.018 - 0.056 | 0.111 | 0.081 | 0.088 | 0.055 - 0.146 |
| validation | 0.101 | 0.093 | 0.076 | 0.040 - 0.128 | 0.125 | 0.105 | 0.104 | 0.055 - 0.157 |
| test | 0.106 | 0.091 | 0.086 | 0.043 - 0.128 | 0.135 | 0.100 | 0.105 | 0.073 - 0.176 |

Table 48: **Absolute error analysis: using two slides together vs. using one of the slides to predict sample-level tumor purity - PRAD cohort.** For a tumor sample with two slides, absolute errors between genomic tumor purity values and sample-level MIL predictions ($e_{smpl}$) and genomic tumor purity values and slide-level MIL predictions ($e_{sld}$) in the training, validation, and test sets are calculated. Mean absolute errors ($\mu_{e_{smpl}}$ and $\mu_{e_{sld}}$) together with standard deviations ($\sigma_{e_{smpl}}$ and $\sigma_{e_{sld}}$) and median absolute errors ($m_{e_{smpl}}$ and $m_{e_{sld}}$) together with interquartile ranges ($IQR_{e_{smpl}}$ and $IQR_{e_{sld}}$) are presented.

| | sample level | | | | slide level | | | |
|---|---|---|---|---|---|---|---|---|
| | $\mu_{e_{smpl}}$ | $\sigma_{e_{smpl}}$ | $m_{e_{smpl}}$ | $IQR_{e_{smpl}}$ | $\mu_{e_{sld}}$ | $\sigma_{e_{sld}}$ | $m_{e_{sld}}$ | $IQR_{e_{sld}}$ |
| training | 0.161 | 0.117 | 0.132 | 0.061 - 0.270 | 0.164 | 0.108 | 0.131 | 0.069 - 0.257 |
| validation | 0.195 | 0.134 | 0.177 | 0.114 - 0.261 | 0.209 | 0.128 | 0.187 | 0.140 - 0.265 |
| test | 0.197 | 0.164 | 0.155 | 0.078 - 0.248 | 0.224 | 0.164 | 0.153 | 0.101 - 0.436 |

Table 49: **Absolute error analysis: using two slides together vs. using one of the slides to predict sample-level tumor purity - UCEC cohort.** For a tumor sample with two slides, absolute errors between genomic tumor purity values and sample-level MIL predictions ($e_{smpl}$) and genomic tumor purity values and slide-level MIL predictions ($e_{sld}$) in the training, validation, and test sets are calculated. Mean absolute errors ($\mu_{e_{smpl}}$ and $\mu_{e_{sld}}$) together with standard deviations ($\sigma_{e_{smpl}}$ and $\sigma_{e_{sld}}$) and median absolute errors ($m_{e_{smpl}}$ and $m_{e_{sld}}$) together with interquartile ranges ($IQR_{e_{smpl}}$ and $IQR_{e_{sld}}$) are presented.

| | sample level | | | | slide level | | | |
|---|---|---|---|---|---|---|---|---|
| | $\mu_{e_{smpl}}$ | $\sigma_{e_{smpl}}$ | $m_{e_{smpl}}$ | $IQR_{e_{smpl}}$ | $\mu_{e_{sld}}$ | $\sigma_{e_{sld}}$ | $m_{e_{sld}}$ | $IQR_{e_{sld}}$ |
| training | 0.087 | 0.105 | 0.064 | 0.027 - 0.097 | 0.096 | 0.108 | 0.062 | 0.033 - 0.108 |
| validation | 0.079 | 0.057 | 0.069 | 0.045 - 0.099 | 0.082 | 0.055 | 0.076 | 0.037 - 0.112 |
| test | 0.115 | 0.130 | 0.083 | 0.032 - 0.133 | 0.119 | 0.123 | 0.081 | 0.063 - 0.115 |

Table 50: **Comparing the absolute errors of sample-level predictions and the expected value of the absolute errors of slide-level predictions in the test sets of different cohorts.** In the test set of each cohort, for a tumor sample with two slides, the absolute errors between genomic tumor purity values and sample-level MIL predictions ($e_{smpl}$) and the expected value of absolute errors between genomic tumor purity values and slide-level MIL predictions ($e_{sld}$) are calculated. Then, the number of samples with two slides (n), the mean absolute errors ($\mu_{e_{smpl}}$ and $\mu_{e_{sld}}$) together with standard deviations ($\sigma_{e_{smpl}}$ and $\sigma_{e_{sld}}$), the median absolute errors ($m_{e_{smpl}}$ and $m_{e_{sld}}$) together with interquartile ranges ($IQR_{e_{smpl}}$ and $IQR_{e_{sld}}$), and the calculated p-values in the statistical tests ($P_{comp}$) are presented. Note that the PRAD (n=21) and UCEC (n=23) cohorts were excluded from this study due to few samples with two slides. The best methods are highlighted in bold.

| | | Sample level | | | | Slide level | | | | |
|---|---|---|---|---|---|---|---|---|---|---|
| | n | $\mu_{e_{smpl}}$ | $\sigma_{e_{smpl}}$ | $m_{e_{smpl}}$ | $IQR_{e_{smpl}}$ | $\mu_{e_{sld}}$ | $\sigma_{e_{sld}}$ | $m_{e_{sld}}$ | $IQR_{e_{sld}}$ | $P_{comp}$ |
| BRCA | 73 | **0.114** | **0.082** | **0.092** | **0.043 - 0.166** | 0.126 | 0.073 | 0.129 | 0.060 - 0.171 | **2.8e-03** |
| GBM | 90 | 0.115 | 0.107 | 0.076 | 0.046 - 0.145 | 0.118 | 0.096 | 0.089 | 0.062 - 0.161 | 7.1e-01 |
| LGG | 31 | 0.178 | 0.149 | 0.146 | 0.100 - 0.218 | 0.168 | 0.152 | 0.106 | 0.067 - 0.198 | 5.6e-01 |
| LUAD | 44 | **0.118** | **0.102** | **0.084** | **0.050 - 0.168** | 0.138 | 0.102 | 0.121 | 0.067 - 0.181 | **3.7e-04** |
| LUSC | 52 | **0.124** | **0.092** | **0.109** | **0.039 - 0.168** | 0.150 | 0.096 | 0.143 | 0.085 - 0.201 | **1.7e-03** |
| OV | 102 | **0.106** | **0.091** | **0.086** | **0.043 - 0.128** | 0.135 | 0.100 | 0.105 | 0.073 - 0.176 | **5.0e-03** |

### 3 Singapore Cohort

Singapore cohort consists of 179 lung adenocarcinoma patients having East Asian ancestry. Each patient has one tumor sample, and one slide is prepared from each tumor sample (except one sample in the training set). The slides are prepared from formalin-fixed paraffin-embedded sections (ffpe).

On the contrary to ffpe sections in the Singapore cohort, slides in the TCGA cohorts are prepared from fresh-frozen sections. These two tissue preservation methods are quite different from each other. While the ffpe method preserves morphology better and is the routine in histopathology, the fresh-frozen method preserves nucleic acids better and is preferred for molecular analysis [4]. Figure 29 presents example patches cropped from the slides of fresh-frozen sections in the TCGA LUAD cohort and ffpe sections in the Singapore LUAD cohort.

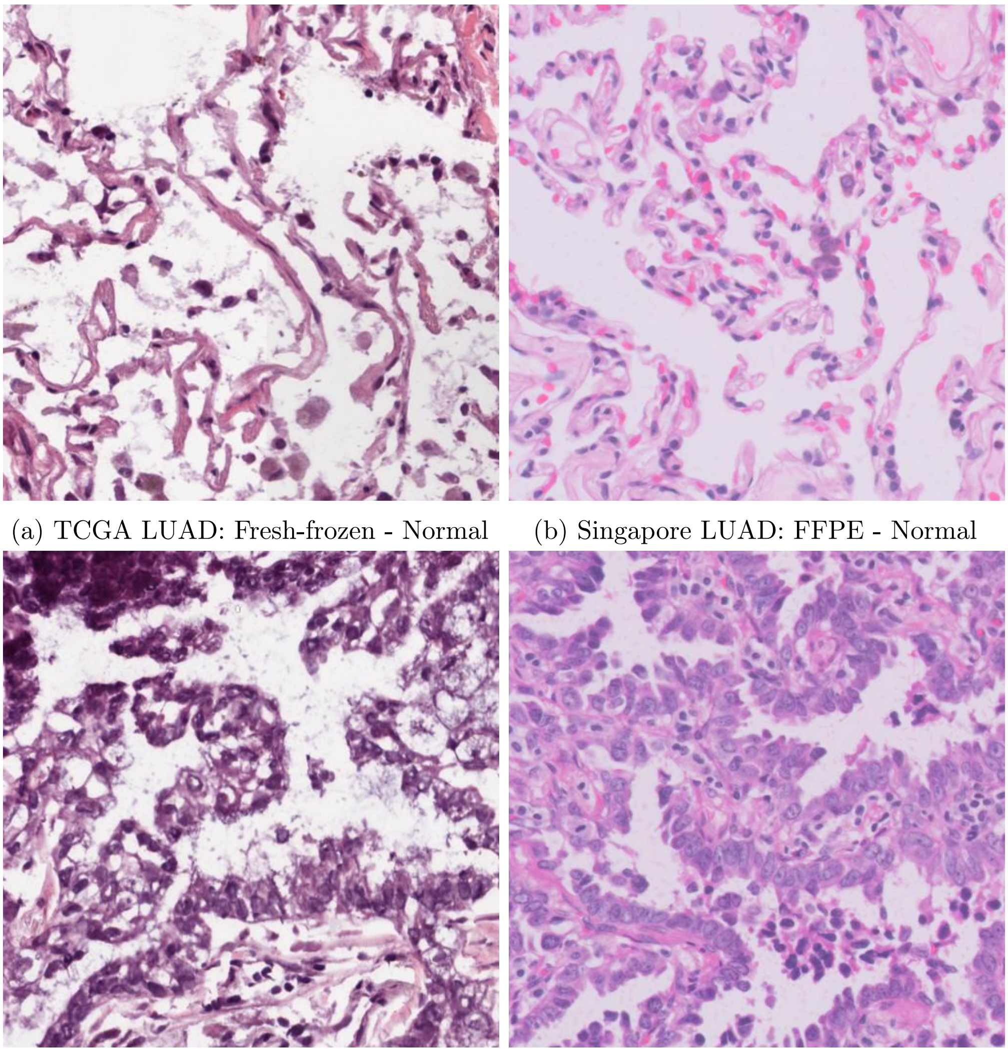

(a) TCGA LUAD: Fresh-frozen - Normal (b) Singapore LUAD: FFPE - Normal

(c) TCGA LUAD: Fresh-frozen - Cancerous (d) Singapore LUAD: FFPE - Cancerous

Figure 29: **Example patches cropped from slides of fresh-frozen and formalin-fixed paraffin-embedded (ffpe) sections.** (**a, c**) A normal patch and a cancerous patch cropped from slides of fresh-frozen sections in the TCGA LUAD cohort. (**b, d**) A normal patch and a cancerous patch cropped from slides of ffpe sections in the Singapore LUAD cohort.

##### 3.1 The number of samples, slides, and patches

The number of samples, slides, and patches in each set are given in Table 51. Note that each patient has only one tumor sample.

Table 51: **Singapore cohort: the number of samples, slides, and patches.**

| | # samples | | | # slides | | | # patches | | |
|---|---|---|---|---|---|---|---|---|---|
| dataset | normal | tumor | total | normal | tumor | total | normal | tumor | total |
| training | 0 | 107 | 107 | 0 | 108 | 108 | 0 | 525,961 | 525,961 |
| validation | 0 | 36 | 36 | 0 | 36 | 36 | 0 | 190,971 | 190,971 |
| test | 0 | 36 | 36 | 0 | 36 | 36 | 0 | 182,383 | 182,383 |
| all | 0 | 179 | 179 | 0 | 180 | 180 | 0 | 899,315 | 899,315 |

##### 3.2 Genomic tumor purity values

We used genomic tumor purity values obtained using ABSOLUTE [1] as labels. Histogram plots of genomic tumor purity values of tumor samples in each set are given in Figure 30.

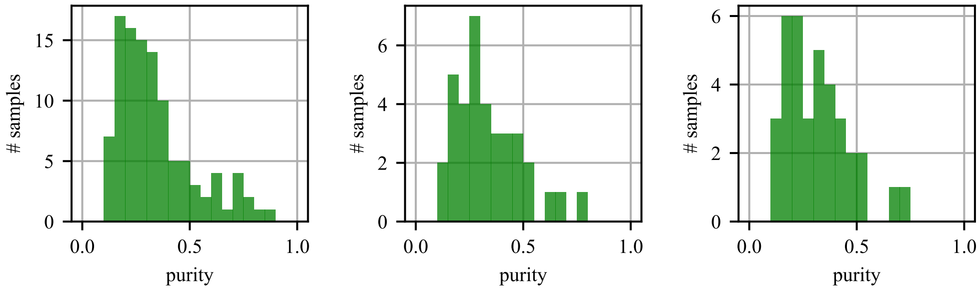


(a) training (b) validation (c) test

Figure 30: **Singapore cohort: genomic tumor purity histograms for training, validation, and test sets.**

##### 3.3 Correlation Analysis

A scatter plot for genomic tumor purity values vs. the MIL model's predictions in each set is given in Figure 31. Moreover, Spearman's correlation coefficients between genomic tumor purity values and the MIL model's predictions in the training, validation, and test sets are presented in Table 52.

##### 3.4 Absolute Error Analysis

The results of absolute error analyses between genomic tumor purity values and the MIL model's predictions in the training, validation, and test sets are presented in Table 53.

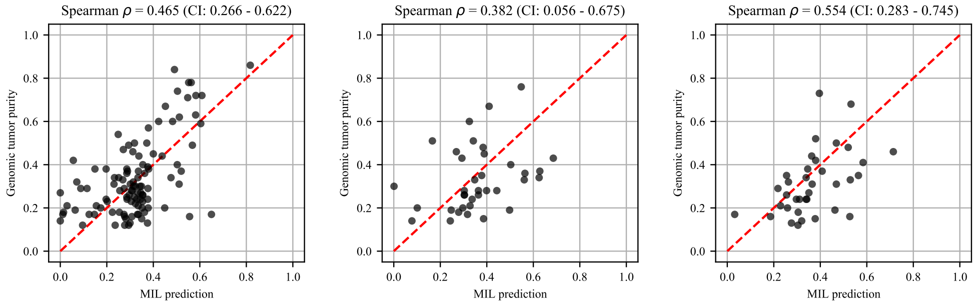


(a) Training (b) Validation (c) Test

Figure 31: **Singapore cohort: scatter plot of genomic tumor purity values and the MIL model's predictions.** Diagonal red dotted lines show the y=x line.

Table 52: **Singapore cohort: correlation analysis.** Spearman's correlation coefficients between genomic tumor purity values and the MIL model's predictions ($\rho_{mil}$) are calculated in the training, validation, and test sets. Correlation coefficients together with calculated p-values ($P_{\rho_{mil}}$) and 95% confidence intervals ($CI_{\rho_{mil}}$) are presented.

| | MIL prediction | | |
|---|---|---|---|
| | $\rho_{mil}$ | $P_{\rho_{mil}}$ | $CI_{\rho_{mil}}$ |
| train | 0.465 | 4.5e-07 | 0.266 - 0.622 |
| valid | 0.382 | 2.1e-02 | 0.056 - 0.675 |
| test | 0.554 | 4.6e-04 | 0.283 - 0.745 |

Table 53: **Singapore cohort: absolute error analysis.** Absolute errors between truth genomic tumor purity values and the MIL model's predictions ($e_{mil}$) in the training, validation, and test sets are calculated. Mean absolute errors ($\mu_{e_{mil}}$) together with standard deviations ($\sigma_{e_{mil}}$) and median absolute errors ($m_{e_{mil}}$) together with interquartile ranges ($IQR_{e_{mil}}$) are presented.

| | MIL prediction | | | |
|---|---|---|---|---|
| | $\mu_{e_{mil}}$ | $\sigma_{e_{mil}}$ | $m_{e_{mil}}$ | $IQR_{e_{mil}}$ |
| training | 0.121 | 0.090 | 0.107 | 0.046 - 0.175 |
| validation | 0.151 | 0.091 | 0.119 | 0.096 - 0.231 |
| test | 0.120 | 0.091 | 0.099 | 0.055 - 0.174 |

### 4 Segmentation of Histopathology Slides in The TCGA LUAD Cohort

In the TCGA LUAD cohort, for each patient with a matching normal sample, we used the trained feature extractor module of our MIL model to extract features of patches cropped over the slides of the tumor and normal samples of the patient. Then, we clustered the patches by using hierarchical clustering over the extracted feature vectors. We determined the distance threshold in hierarchical clustering such that there were 4 clusters among the patches from slides of the normal sample. This made our clustering approach robust against patient-to-patient variations. Indeed, this was why we decided to use both tumor and normal samples of the patient. In other words, instead of determining a global distance threshold for all patients, we calculated patient-specific distance threshold values to capture inter-patient variations.

Each cluster can be assigned one of two labels: cancerous or normal. Ideally, a cluster with a cancerous label can contain patches only from slides of the tumor sample. On the other hand, a cluster with a normal label can contain patches from slides of both the tumor and the normal samples since the tumor sample may also contain normal tissue components. As a post-processing step, we analyzed normal clusters. If the number of patches from slides of the normal sample in a normal cluster was less than 10%, we split this cluster into two such that patches from slides of the tumor sample were assigned to a new cancerous cluster. Finally, we created segmentation masks for slides of the tumor sample by using cluster labels assigned to the patches.
